# Sample preparation and imaging conditions affect mEos3.2 photophysics in fission yeast cells

**DOI:** 10.1101/2020.05.28.119735

**Authors:** Mengyuan Sun, Kevin Hu, Joerg Bewersdorf, Thomas D. Pollard

## Abstract

Photoconvertible fluorescent proteins (PCFPs) are widely used in super-resolution microscopy and studies of cellular dynamics. However, our understanding of their photophysics is still limited, hampering their quantitative application. For example, we do not know the optimal sample preparation methods or imaging conditions to count protein molecules fused to PCFPs by single-molecule localization microscopy in live and fixed cells. We also do not know how the behavior of PCFPs in live cells compares with fixed cells. Therefore, we investigated how formaldehyde fixation influences the photophysical properties of the popular green-to-red PCFP mEos3.2 in fission yeast cells under a wide range of imaging conditions. We estimated photophysical parameters by fitting a 3-state model of photoconversion and photobleaching to the time course of fluorescence signal per yeast cell expressing mEos3.2. We discovered that formaldehyde fixation makes the fluorescence signal, photoconversion rate and photobleaching rate of mEos3.2 sensitive to the buffer conditions by permeabilizing the yeast cell membrane. Under some imaging conditions, the time-integrated mEos3.2 signal per yeast cell is similar in live cells and fixed cells imaged in buffer at pH 8.5 with 1 mM DTT, indicating that light chemical fixation does not destroy mEos3.2 molecules. We also discovered that some red-state mEos3.2 molecules entered an intermediate dark state that is converted back to the red fluorescent state by 561-nm illumination. Our findings provide a guide to compare quantitatively conditions for imaging and counting of mEos3.2-tagged molecules in yeast cells. Our imaging assay and mathematical model are easy to implement and provide a simple quantitative approach to measure the time-integrated signal and the photoconversion and photobleaching rates of fluorescent proteins in cells.

**STATEMENT OF SIGNIFICANCE:** Making quantitative measurements with single-molecule localization microscopy (SMLM) has been impeded by limited understanding of the photophysics of the fluorophores, which is very sensitive to the sample preparation and imaging conditions. We characterized the photophysics of the green-to-red photoconvertible fluorescent protein mEos3.2, which is widely used in SMLM. We combined quantitative fluorescence microscopy and mathematical modeling to measure the fluorescence signal and rate constants for photoconversion and photobleaching of mEos3.2 in live and fixed cells under a wide range of illumination intensities. Our findings provide a guide to compare conditions for imaging and counting mEos3.2-tagged proteins in cells. The presented approach is generally applicable to characterize other fluorescent proteins or dyes in cells.

## INTRODUCTION

Photoactivatable or photoconvertible fluorescent proteins (PAFPs or PCFPs) have enabled super-resolution imaging by temporally separating closely-spaced molecules (1–3). The fluorescent protein EosFP (4) and its derivatives (5–7) have been widely used in SMLM for both live (8–10) and fixed biological samples (11–13). The fluorescent protein is fused to the coding sequence of a protein of interest in the genome for endogenous expression or expressed exogenously and transiently in cells. Irradiation at 405 nm photoconverts EosFPs from their native green state with an emission peak at 516 nm to their red state with an emission peak at 580 nm (4, 14, 15). In SMLM, sparsely distributed photoconverted red EosFPs are excited at 561 nm and then localized (13). The rationally designed, monomeric derivative of EosFP mEos3.2 is favored by many due to its monomeric property, high brightness, photostability, and compatibility with live cells (6).

Counting fluorescently-tagged fusion proteins is a potential strength of SMLM, as the images are assembled from discrete localizations of individual molecules (16, 17). The total number of localizations in the SMLM images is closely correlated to the total number of fusion proteins, which allows the measurement of this important quantity even in a diffraction-limited subcellular structure (9, 11, 12, 18–23). Genetically encoded tagging with PAFPs or PCFPs can ensure 1:1 labeling stoichiometry (16), without the uncertainties associated with extrinsic labeling techniques (22, 24–28). However, even with genetically encoded tags, quantitative SMLM still faces several challenges that can lead to undercounting or overcounting the molecules. Fluorescent proteins mature slowly, so an unknown fraction of the FPs is fluorescent at the time of imaging (29). Some of the PAFPs or PCFPs might never be photoconverted or photoactivated to the active state for SMLM imaging (30). Moreover, activated PAFPs or PCFPs may blink by entering a transient dark state and return to the fluorescent state an unknown number of times, which can lead to overcounting (18, 20). A promising way of doing quantitative SMLM is to obtain numbers of the tagged molecules relative to internal calibration standards of known number (9, 11, 12, 21, 23). The sources of error mentioned above can be accounted for if the target and calibration standards are prepared, imaged, and analyzed consistently in the same way (11, 12, 16, 31).

However, any inconsistency in the process of sample preparation, SMLM imaging, or data analysis can introduce errors. Diffusion and other movements of the fluorescent protein can further complicate the quantification process, so light chemical fixation can be used to preserve the targeted structure and eliminate movements for quantitative SMLM of cells expressing proteins tagged with mEos2 (11, 12). However, fixation can introduce measurement errors. For example, fixation might destroy some FPs or change their photophysical properties (32), which can change the average number of localizations for the FP. Therefore, one must understand how fixation and sample preparation affect the mEos3.2 photophysical parameters related to single-molecule imaging and counting. Photoconversion and photobleaching rates determine the density of active fluorophores at each frame, which is essential for separating closely-spaced molecules. The fluorescence signal of fluorescence fusion proteins in the structure of interest contains information about the brightness of individual molecules and the number of molecules able to fluoresce, both aspects being important for molecule counting with diffraction-limited (31) and super-resolution imaging (17).

In this study, we measured the photophysical properties of mEos3.2 in fission yeast cells by fitting a 3-state model of photoconversion and photobleaching to the time course of the mEos3.2 fluorescence signal per cell measured by quantitative fluorescence microscopy. We found that formaldehyde fixation permeabilized the *S. pombe* cells for small molecules, making the photophysical properties of mEos3.2 sensitive to the buffer conditions. To find conditions where the mEos3.2 photophysical parameters are comparable in live and fixed yeast cells, we investigated the effect of fixation and imaging buffer under a wide range of imaging conditions with point-scanning and widefield illumination. We also discovered that a subpopulation of red-state mEos3.2 molecules entered an intermediate state that is converted back to the red fluorescent state by illumination at 561-nm but not 405-nm. Our data provide information on sample preparation for imaging and counting mEos3.2 in live and fixed yeast cells. Our quantitative imaging assay combined with the 3-state model can be applied to study the photophysical properties of other PAFPs and PCFPs quantitatively without doing single-molecule localization.

## MATERIALS AND METHODS

### Plasmids and *S. pombe* Strains

The open reading frame encoding mEos3.2 was cloned into the pJK148-pAct1-nmt1Term plasmid with PCR and NEB HiFi Builder Assembly. Both the newly constructed plasmid and chromosomal insertion were verified by sequencing.

### Preparation of *S. pombe* Cells for Imaging

*S. pombe* cells expressing mEos3.2 were grown in exponential phase at 25 °C in YE5S-rich liquid medium in 50-mL flasks in the dark before switching to EMM5S-synthetic medium ~12-18 hours before imaging to reduce the cellular autofluorescence background. Live cells were concentrated 10- to 20-fold by centrifugation at 3,000 rpm for 30 s and resuspended in EMM5S for imaging.

Cells were fixed by mixing an equal volume of fresh, room temperature 4% formaldehyde aqueous solution (EMS) with the cell culture and shaking at 150 rpm at 25° C for 15 min or 30 min. The cells were pelleted by centrifugation at 3,000 rpm for 30 s and washed by pelleting in EMM5S or other buffers 3 times, and then resuspended in EMM5S or other buffers. Concentrated cells in 5 μL were mounted on a thin layer formed from 35 μL of 25% gelatin (Sigma-Aldrich; G-2500) in EMM5S or other buffers (without the DTT). To assess how pH and reducing agent affect the photophysical properties of mEos3.2, we fixed cells for 30 min and then washed and resuspended the cells for imaging in one of the following buffers: 50 mM MES (pH 5.5), 50 mM MES (pH 6.5), 50 mM Tris-HCl (pH 7.5), 50 mM Tris-HCl (pH 8.5), 50 mM Tris-HCl (pH 7.5) with 1 mM DTT, 50 mM Tris-HCl (pH 8.5) with 1 mM DTT, or 50 mM Tris-HCl (pH 8.5) with 10 mM DTT.

### Point-scanning Confocal Imaging Conditions

Time lapse videos were acquired on a Zeiss Laser Scanning Microscope (LSM 880) using an alphaPlan-Apochromat 100x/NA 1.46 oil-immersion objective and an emission band path filter collecting fluorescence in the 566 - 719 nm wavelength range. Samples were illuminated by scanning a field of view (FOV) of 85 x 85 μm (512 x 512 pixels; 160 nm pixel size) with both the 405 nm and 561 nm lasers at constant intensities. To test different imaging conditions, we set the 405 nm laser power at the sample constant ranging from 16 to 56 μW and the 561 nm laser power constant ranging from 11 to 37 μW. To compare with widefield imaging conditions, we estimated the average intensity and the peak intensity of point-scanning illumination. The average intensities in the FOV (power at the sample divided by the FOV area) were 0.22 to 0.78 W/cm^2^ at 405 nm and 0.15 to 0.51 W/cm^2^ at 561 nm. The peak intensities (power at the sample divided by the size of the point spread function) were ~80 to 240 kW/cm^2^ at 405 nm and ~20 to 80 kW/cm^2^ at 561 nm. A Z-stack of 19 slices spaced at 600-nm intervals was acquired with a pixel dwell time of 0.85 μs. The total exposure time for each Z-stack was 4.23 s (0.85 μs x 512 x 512 x 19). An entire time lapse data set consisted of 50 or 100 Z-stacks at an acquisition rate of approximately 4 Z-stacks per minute (due to overhead in the scan process).

For experiments with alternating 405 and 561-nm laser illumination, the 561-nm laser scanned the FOV for 10 cycles followed by 405-nm laser illumination for 5 cycles with either no break or a 2 min break between the 405-nm period and the following 561-nm illumination period. The laser powers at the sample were 56 μW at 405 nm and 37 μW at 561 nm. This procedure was repeated 7 times. We increased the temporal resolution by reducing the pixel number in the 85 x 85 μm FOV to 128 x 128 (640 nm pixel size) with a pixel dwell time of 3.39 μs. This approach reduced the total exposure time for each Z-stack to 1.05 s (3.39 μs x 128 x 128 x 19). These 3D stacks were collected with 40 cycles of 561-nm illumination followed by 20 cycles of 405-nm irradiation without breaks between the 405-nm and 561-nm illumination periods. The Z-stacks were acquired at a rate of approximately 5 Z-stacks per minute due to the scanning overhead. This procedure was repeated 7 times.

For GFP photobleaching measurements, wildtype cells and cells expressing Fim1-GFP (TP347) were excited at 488 nm (~60 μW at the sample) and emission fluorescence in the range of 505-735 nm was collected. A 19-slice Z stack with 600-nm Z-step intervals covering a FOV of 82 x 82 μm (512 x 512 pixels; 160 nm pixel size) was imaged at each time point with a pixel dwell time of 0.85 μs. The entire time-lapse data set consisted of 100 Z-stacks.

### Widefield Fluorescence Imaging Conditions

Time lapse videos were acquired with a custom-built single-molecule localization microscope (SMLM) based on a Leica DMi8 stand with widefield illumination, a 63x/1.47 NA oil-immersion objective, and a band pass filter to collect emission fluorescence in the 584-676 nm wavelength range. Samples were illuminated at both 405 nm and 561 nm and imaged with an sCMOS camera (Hamamatsu ORCA-Flash4.0 V2) at 50 frames per second (fps) for 15,000 frames. To test different imaging conditions, the 405-nm laser intensity was set constant in a range from 0.5 to 2 W/cm^2^, and the 561-nm laser intensity from 1 W/cm^2^ to 1 kW/cm^2^. A custom-written LABVIEW program was used to control the lasers for alternating 405-nm and 561-nm illumination.

### Image Analysis

Images recorded by confocal and widefield fluorescence microscopes were viewed and analyzed in Fiji (Fiji is Just ImageJ) (33). We made a sum projection of the 19-slice Z-stacks of the time-lapse confocal images. We manually selected a region of interest (ROI) 1 (containing typically ~50-100 cells for the confocal images and ~ 25 cells for the widefield images) with the polygon tool and selected the background ROI 2 with the square tool (Fig. S1). The area and mean signal per pixel (MSPP) of both ROIs were measured and the fluorescence signal per cell at each time point was calculated based on: [Area 1 * (MSPP 1 – MSPP 2)] / number of cells in ROI 1. We calculated the weighted mean and standard deviation of the fluorescence signal per cell from all the FOVs included for each condition, weighted by the number of cells in each FOV. To correct for autofluorescence background, we subtracted the autofluorescence signal per wildtype cell at each time point from the fluorescence signal per cell expressing mEos3.2.

### Analytical Model of the Time Course of Red mEos3.2 Fluorescence Signal per Cell

Our three-state model (Fig. 1C) considers mEos3.2 molecules to have 3 different states: a non-activated green (G) state, an activated red (R) state, and a bleached (B) state. Photoconversion converts molecules from the G- to the R-state by an irreversible first-order reaction with a rate constant of k_act_ (k_activation_). Molecules in the R-state emit red photons until is photobleached to the B state by an irreversible first-order reaction with a rate constant of k_bl_ (k_bleaching_). With illumination at 405 nm and 561 nm, the rates of change in the numbers (n) of G-, R-, and B-state mEos3.2 molecules are described by the following differential equations (Fig. S2A):

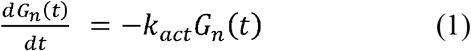

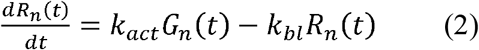

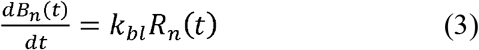

**Figure 1:**
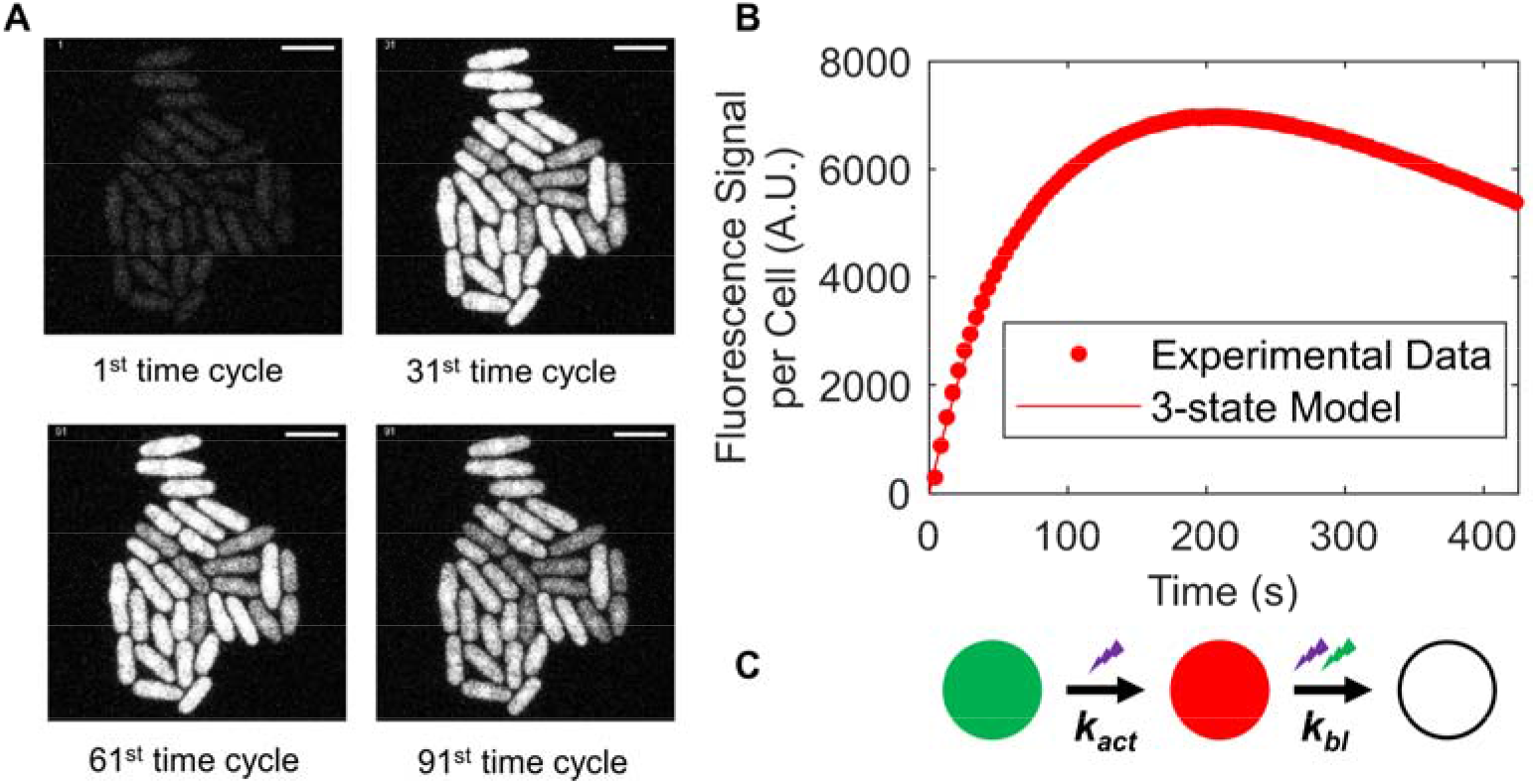
Photoconversion and photobleaching of mEos3.2. **(A)** Time series of fluorescence micrographs of a field of *S.pombe* cells expressing cytoplasmic mEos3.2 at the 1^st^, 31^st^, 61^st^, and 91^st^ time cycles. At each of the 100 cycles, a point-scanning confocal microscope illuminated the cells simultaneously at both 405 nm and 561 nm, 19-slices in a Z stack were imaged with a total exposure time of 4.23 s and sum-projected with the same contrast. Scale bar = 10 μm. **(B)** Time course of the fluorescence signal per cell at 566-719 nm (after autofluorescence subtraction). Fitting Eq. 7 of the 3-state model in panel C (line) to the data (dots) gave a photoconversion rate constant (k_act_) of 1.2 x 10^-2^ s^-1^ (95% CI: 1.16 −1.24 x 10^-2^) and a photobleaching rate constant (k_bl_) of 1.6 x 10^-3^ s^-1^ (95% CI: 1.5-1.7 x10^-3^). **(C)** Three state model for mEos3.2 photoconversion and bleaching. Illumination at 405 nm photoconverts mature mEos3.2 molecules from the green (G) state to the red (R) state with a photoconversion rate constant of k_act_. Illumination at both wavelengths photobleaches red mEos3.2 molecules with a rate constant of k_bl_.

We defined the total number of mEos3.2 molecules in a cell as M_n_, and assumed that all mEos3.2 molecules were in the green state at the start of the experiment, so G_n_(t = 0) = M_n_, R_n_(t = 0) = 0, and B_n_(t = 0) = 0. Solving the system of differential equations analytically resulted in the following equations for the number of G-, R-, and B-state molecules in a cell changing over time:

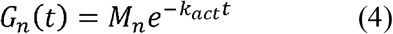

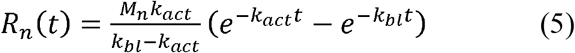

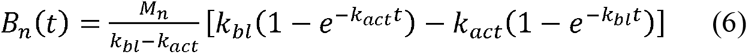

Eq. 5 describes how the number of R-state mEos3.2 molecules in a cell (R_n_) changes with continuous photoconversion and photobleaching. Assuming that the signal of an R-state molecule per frame is ε_f_, the fluorescence signal from the red mEos3.2 molecules per cell (R_s_) at each frame recorded at a given time t is R_s_(t) = R_n_(t) x ε_f_. Multiplying both sides of Eq. 5 by ε_f_ gives Eq. 7 that describes how the fluorescence signal per cell in each frame R_s_(t) changes over time with continuous photoconversion and photobleaching:

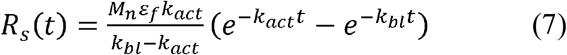

We estimated M_n_ x ε_f_, k_act_, and k_bl_ using Levenberg-Marquardt nonlinear least squares regression to fit Eq. 7 to the time course of fluorescence signal per cell R_s_(t). We calculated the 95% confidence intervals (CI) of the fitted parameters from the covariance argument of the fit. For the confocal experiments, we fit Eq. 7 to the weighted average time course of fluorescence signal per cell from all FOVs and report the 95% CI of the fitted parameters. For the widefield illumination experiments, we fit Eq. 7 individually to the time course of fluorescence signal per cell (after autofluorescence background subtraction) from each FOV. We calculated the mean fitted parameters and standard deviation weighted by the number of cells in each FOV.

Since R_s_(t) is the fluorescence signal from red mEos3.2 molecules per cell in each frame, the fluorescence signal from red mEos3.2 molecules per cell per second is R_s_(t) x f (frame rate, fps). Integrating the function R_s_(t) x f with respect to time (t, second) over the interval of [0, ∞] gives the time-integrated signal 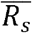 of mEos3.2 per cell:

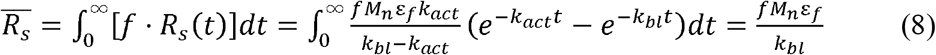

We used Eq. 8 to calculate 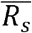 using the parameters M_n_ x ε_f_ and k_bl_ from the previous fit of Eq. 7. We estimated the 95% CI or standard deviation through error propagation.

Experiments with alternating illumination at 405 nm and 561 nm revealed a fourth intermediate (I) state of mEos3.2 molecules (Fig. 6). Illumination at 405 nm converts mEos3.2 molecules in the G-state into the R-state and molecules in the G- and/or R-state into the I-state. Irradiation at 561 nm converts the mEos3.2 molecules from the I-state to R-state with an activation rate constant of k_act,561_. R-state molecules are photobleached with a bleaching rate constant of k_bl_. During a 561-nm illumination period after a previous 405-nm illumination period, we assumed that no mEos3.2 molecules in the G- and/or R-state converted to the I-state and I-states molecules were not photobleached. The rates of change in the numbers (n) of I-, R-, and B-state mEos3.2 molecules are described by the following differential equations:

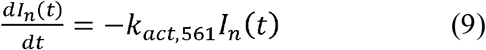

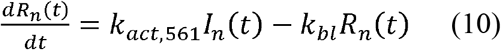

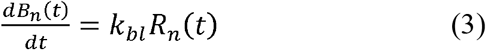

We defined the total number of mEos3.2 molecules, i.e. the sum of molecules in the I-, R- and B-states in a cell, after the previous 405-nm irradiation as S_n_. We further defined t = 0 as the time at which 561-nm illumination starts and assumed that S_n_ is constant during the 561-nm illumination period, since the 405-nm laser was off and conversion of molecules to the I-state by 561-nm light is negligible. We further assume that the number of molecules in the different states at the beginning of the 561-nm illumination period is R_n_(t = 0) = R_n,0_, I_n_(t = 0) = S_n_ - R_n,0_, and B_n_(t = 0) = 0. Solving the system of differential equations analytically resulted in the following equations for the number of I-, R-, B-state molecules changing over time:

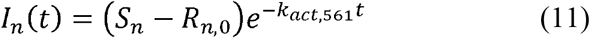

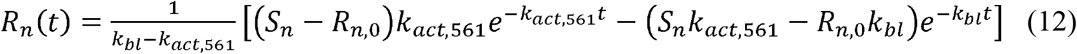

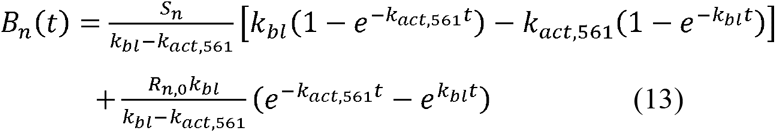

Eq. 12 describes how the number of red mEos3.2 molecules in a cell (R_n_) changes during the period of 561-nm illumination. Multiplying both sides of Eq. 12 by ε_f_ gives equation (14) that describes how the fluorescence signal per cell changes over time during this period:

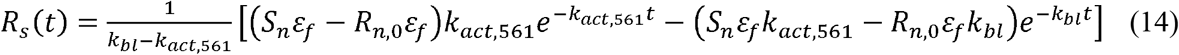

We estimated S_n_ x ε_f_, R_n,0_ x ε_f_, k_act,561_, and k_bl_ using Levenberg-Marquardt nonlinear least squares regression to fit Eq. 14 to the time courses of the fluorescence signal per cell during the 561-nm illumination period. We calculated the mean and standard deviation of the activation rate constant (k_act,561_) for converting mEos3.2 molecules from I- to R-state by 561-nm irradiation from averaging the 7 periods of 561-nm irradiation for each condition.

### Single-molecule Localization

We acquired data on single R-state mEos3.2 molecules with our custom-built SMLM using a 405-nm laser intensity of 1 W/cm^2^, a 561-nm laser intensity of 1 kW/cm^2^, and a frame rate of 50 fps. We localized single molecules with the Python Microscopy Environment (PYME) package (34), using a threshold of 0.6 for event detection computed from the estimated pixel signal-to-noise ratio (SNR). We corrected pixel-dependent noise with maps generated from dark camera frames. We measured the number of photons from single R-state mEos3.2 molecules in each 20-ms frame between frames 5,000 and 10,000 only, since the R-state molecules before frame 5,000 were too dense for localization and most were bleached after frame 10,000.

### Cell Permeability Experiments

Wildtype *S. pombe* cells were fixed for 15 min or 30 min as above. Fixed and live cells were mounted on coverslips coated with 0.5 mg/mL peanut-lectin (Sigma-Aldrich; L0881-5MG) for 30 min. The cells were then incubated for 30 min in EMM5S medium containing 20 μM fluorescein (Sigma-Aldrich 46960-25G-F) or fluorescein-dextran 3,000 MW (ThermoFisher D3305). The live cells mixed with the dyes were imaged at room temperature (~23° C) immediately to minimize endocytosis of the dyes. Wildtype cells in EMM5S medium alone were prepared and imaged as negative controls for autofluorescence subtraction. Brightfield and confocal fluorescence images of the mid-sections of cells were acquired with LSM880. The samples were excited at 488 nm (~22 μW at the sample) and the emitted fluorescence was collected using an emission band path filter with a 519-601 nm wavelength range. We analyzed the images in Fiji (33) by manually selecting ROIs of 5×5-pixel squares in the cells (~30-40 cells for each condition; 1 ROI per cell) and the extracellular environment (6~12 ROIs for each condition) (Fig. S3A). We measured the MSPP of all ROIs including the negative controls (for autofluorescence subtraction), quantified the cell permeability in each condition by calculating (MSPP_cytoplasmic_ – MSPP_negative control_)/ MSPP_extracellular_, and reported the mean and standard deviation.

## RESULTS

### Quantitative assessment of mEos3.2 photophysics in yeast cells by fitting a 3-state model to fluorescence microscopy data

We combined quantitative fluorescence microscopy with mathematical modeling to estimate the time-integrated signal per cell detected in the red channel 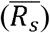, and the rate constants for photoconversion and photobleaching (k_act_ and k_bl_) of mEos3.2 in the cytoplasm of fission yeast cells (Fig. 1). Our model assumes that illumination at 405 nm photoconverts mEos3.2 molecules irreversibly from their green (G) to their red (R) state with an activation rate constant of k_act_ and that the 561-nm light excites the red-form mEos3.2 with a peak emission at ~580 nm. Illumination at either 405 nm or 561 nm converts R-state mEos3.2 molecules to the bleached (B) state by an irreversible first-order reaction with a bleaching rate constant of k_bl_ (Fig. 1C). Our 3-state model did not consider mEos3.2 photoswitching or “blinking” in its G- or R-state, where the protein enters a transient off state and can be converted back to the fluorescent state.

Fission yeast cells expressing mEos3.2 from the actin promoter in the *leu1* locus ensured a relatively high and homogenous cytoplasmic mEos3.2 expression level (Fig. 1A). We used point-scanning confocal microscopy to illuminate the cells at both 405 nm and 561 nm and collect time-lapse images in the red wavelength range of 566-719 nm (Fig. 1A). The time course of the fluorescence signal per cell first rose as the large pool of molecules in the G-state was photoconverted to the R-state, from which we detected red photons as signal, and then declined as the pool of molecules in the G-state was depleted and photobleaching depleted the R-state pool (Fig. 1B).

The equation (Eq. 7) of our 3-state model (Fig. 1C) fit the time courses of fluorescence signal per cell very closely (Fig. 1B). The best fits yielded estimates of the product of total number of molecules per cell and detected signal per R-state molecule per frame (M_n_ x ε_f_), and the rate constants for photoconversion (k_act_) and photobleaching (k_bl_) (Table S1). We then used Eq. 8 to calculate the time-integrated signal per cell 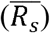 using fitted parameters M_n_ x ε_f_ and k_bl_ (Table S1). We used this approach to measure how sample preparation and imaging conditions influence these photophysical properties.

### Effects of formaldehyde fixation and imaging buffer on photophysical properties of mEos3.2 in fission yeast cells

We investigated how fixation affects the photophysical properties of mEos3.2, so we could compare experiments on live and fixed yeast cells (Fig. 2). Yeast cells fixed with formaldehyde in EMM5S-synthetic growth medium emitted fewer red photons than live cells imaged under the same conditions (Fig. 2C). Fitting Eq. 7 of the 3-state model (Fig. 1C) to the time courses of fluorescence signal per yeast cell showed that mEos3.2 in fixed cells had lower M_n_ x ε_f_ (Fig. 2D) and 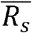 (Fig. 2G) and higher k_act_ (Fig. 2E) and k_bl_ (Fig. 2F) than live cells. Yeast cells fixed for 30 min had even lower fluorescence signals (Fig. 2D, G) and higher rate constants than cells fixed for 15 min (Fig. 2E, F).

**Figure 2:**
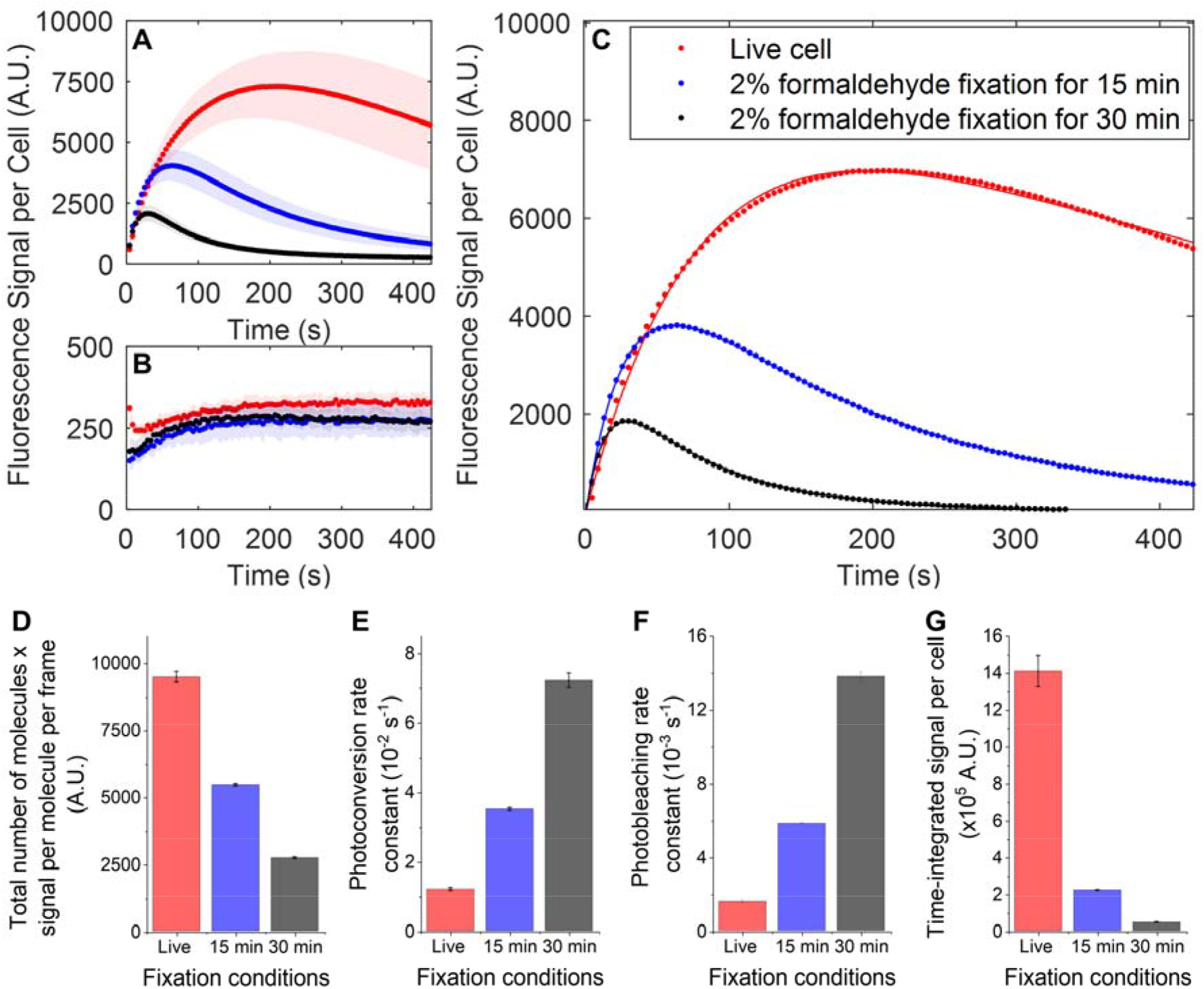
Effects of formaldehyde fixation on the fluorescence signal of mEos3.2 and rate constants for photoconversion and photobleaching. **(A**) Time courses of the fluorescence signal per *S. pombe* cell expressing cytoplasmic mEos3.2 illuminated at 405 nm (22 μW) and 561 nm (15 μW) by point-scanning confocal microscopy under 3 conditions: live cells (red dots) or cells fixed with 2% formaldehyde for 15 min (blue dots) or 30 min (black dots) in EMM5S medium and imaged in EMM5S. Nine fields of view (FOV) of 85 μm x 85 μm were taken over time for each condition. Plots are weighted mean (dots) and standard deviations (shaded area) of the fluorescence signal per cell. **(B)** Time courses of the total autofluorescence signal per wildtype *S. pombe* cell under the same conditions as in panel A. Four FOVs of 85 μm x 85 μm over time were taken for each condition. Plots show weighted mean (dots) and standard deviations (shaded area) of the fluorescence signal per cell. **(C)** Time courses of the mEos3.2 fluorescence signal per cell at 566-719 nm after autofluorescence subtraction. Eq. 7 of the 3-state model was fit to the experimental data (dots). The lines are theoretical curves using the parameters that best fit the data. **(D-G)** Comparison of parameters of live cells and cells fixed for 15 or 30 min. The error bars are 95% confidence intervals of the parameters (Table S1). **(D)** The product of total number of molecules per cell and signal of an R-state mEos3.2 molecule per frame (M_n_ x ε_f_) from the fit. **(E)** Photoconversion rate constant (k_act_) from the fit. **(F)** Photobleaching rate constant (k_bl_) from the fit. **(G)** Time-integrated signal per cell 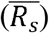 calculated using Eq. 8.

We used fluorescein to test our hypothesis that formaldehyde fixation affects mEos3.2 photophysics by permeabilizing the yeast cell membrane for small molecules (Fig. S3). Live yeast cells excluded fluorescein (332 g/mol), but fixation with formaldehyde (without detergents or organic solvents) partially permeabilized the cell membrane, allowing the entry of fluorescein (Fig. S3D). Moreover, more fluorescein entered the cells fixed for 30 min than the cells fixed for 15 min (Fig. S3D). Thus, ions and small molecules in the imaging buffer, such as thiol DTT (154 g/mol), equilibrated with the interior of the fixed yeast cells and affected the photophysical properties of mEos3.2.

Knowing that fixed *S. pombe* cells are permeable for small molecules, we tested how the composition of the imaging buffer influenced mEos3.2 photophysics (Fig. 3). We hypothesized that the photophysical changes of mEos3.2 in the fixed cells were due to the low pH (~5.5) and oxidizing environment of EMM5S relative to the live-cell cytoplasmic environment. We found that 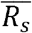 was higher (Fig. 3B), and k_act_ (Fig. 3D) and k_bl_ (Fig. 3E) were lower in imaging buffers with higher pH. Adding the reducing agent DTT to the imaging buffer further increased 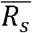 (Fig. 3B) and decreased k_bl_ in the pH range we tested (Fig. 3E). A concentration of 1 mM DTT was more effective than 10 mM DTT at increasing 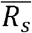 (Fig. 3B). Additionally, the value of M_n_ x ε_f_ from fixed cells in 50 mM Tris-HCl (pH 8.5) with 1 mM DTT (9187 A.U., 95% CI: 9024 – 9351, Table S2) was similar to that from live cells (9489 A.U., 95% CI: 9291 - 9687, Table S1). Values of 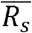 were also similar in live cells and fixed cells at pH 8.5 with 1 mM DTT (Table S1, S2). Thus, mEos3.2 molecules survived fixation and the total number of fluorescence-competent molecules per cell (M_n_) did not change, but the extracellular imaging buffer influenced the intracellular mEos3.2 signal per frame (ε_f_) and other photophysical properties as photoconversion and photobleaching rates. In the following, we therefore used the imaging buffer at pH 8.5 with 1 mM DTT (called ‘Tris8.5-DTT buffer’) for many fixed cell imaging conditions.

**Figure 3:**
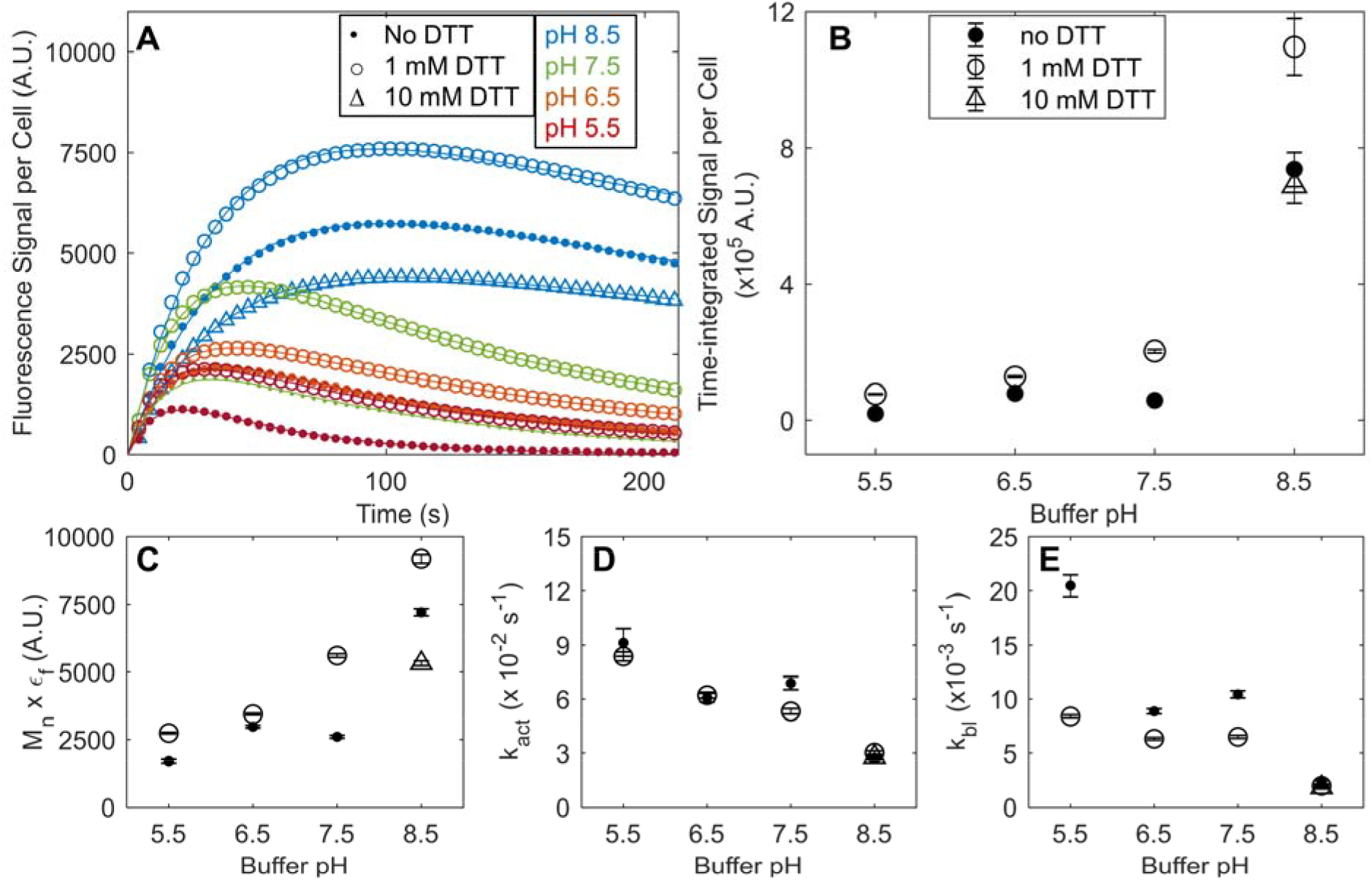
Effects of pH and DTT on the fluorescence signal and rate constants for photoconversion and photobleaching of mEos3.2 in fixed *S. pombe* cells. **(A)** Time courses of the fluorescence signal per cell expressing cytoplasmic mEos3.2 (after autofluorescence subtraction). The cells were fixed with 2% formaldehyde for 30 min and illuminated at 405 nm (22 μW) and 561 nm (15 μW) by point-scanning confocal microscope under 9 different buffer conditions: (red dot) 50 mM MES (pH 5.5); (red circle) 50 mM MES (pH 5.5) with 1 mM DTT; (orange dot) 50 mM MES (pH 6.5); (orange circle) 50 mM MES (pH 6.5) with 1 mM DTT; (green dot) 50 mM Tris-HCl (pH 7.5); (green circle) 50 mM Tris-HCl (pH 7.5) with 1 mM DTT; (blue dot) 50 mM Tris-HCl (pH 8.5); (blue circle) 50 mM Tris-HCl (pH 8.5) with 1 mM DTT: (blue triangle) 50 mM Tris-HCl (pH 8.5) with 10 mM DTT. The continuous lines are best fits of Eq. 7 of the 3-state model to the time courses of the fluorescence signal per cell. Fig. S7 reports the raw data. **(B-E)** Dependence of the parameters on pH and DTT. The error bars are 95% confidence intervals for the parameters (Table S2). **(B)** Time-integrated signal per cell 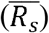 calculated using Eq. 8. **(C)** The product of total number of molecules per cell and the signal of an R-state mEos3.2 molecule per frame (M_n_ x ε_f_) from the fit. **(D)** Photoconversion rate constant (k_act_) from the fit. **(E)** Photobleaching rate constant (k_bl_) from the fit.

### Effects of the 405-nm and 561-nm laser intensities on photophysical properties of mEos3.2 in fission yeast cells

We used laser-scanning confocal microscopy and widefield fluorescence microscopy to test the effects of a wide range of laser intensities on mEos3.2 photophysics in fixed *S. pombe* cells in Tris8.5-DTT buffer (Fig. 4). The products of total mEos3.2 molecules per cell and signal per R-state molecule per frame (M_n_ x ε_f_) were similar in live cells and fixed cells in Tris8.5-DTT buffer (Fig. 4A-D). This was true for low-power laser-scanning confocal microscopy conditions as well as widefield SMLM imaging conditions with a 405-nm laser intensity of 0.5-2 W/cm^2^ and a 561-nm laser intensity of 1 kW/cm^2^ (Fig. 4C). However, at widefield 561-nm laser intensities of 10 and 100 W/cm^2^, the time-integrated signal per cell 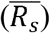 differed in live cells and fixed cells in Tris8.5-DTT buffer (Fig. 4P). Fixation and the imaging buffer had different effects on M_n_ x ε_f_ (Fig. 4D), k_act_ (Fig. 4H), k_bl_ (Fig. 4L) and 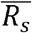 (Fig. 4P) of mEos3.2 depending on the 561-nm laser intensities.

**Figure 4:**
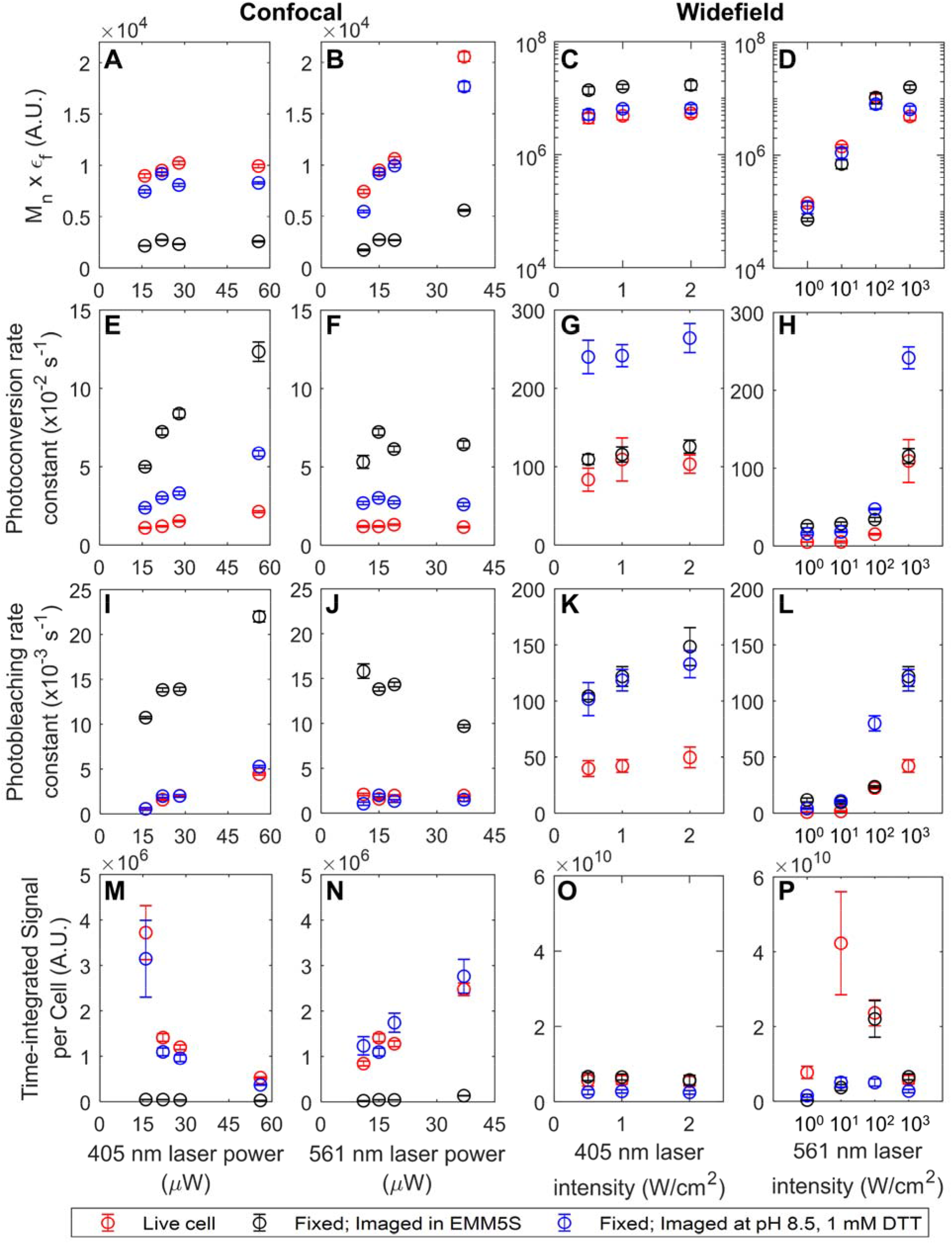
The effects of 405-nm and 561-nm laser intensities on the fluorescence signal and rate constants for photoconversion and photobleaching of mEos3.2 in live and fixed *S. pombe* cells. Cells expressing cytoplasmic mEos3.2 were imaged live (red circles) or fixed with 2% formaldehyde for 30 min in EMMS5 medium and mounted in EMM5S medium (black circles) or 50 mM Tris-HCl (pH 8.5) with 1 mM DTT (blue circles). Cells were imaged with 7 different laser intensities by confocal microscope and 6 different laser intensities by widefield microscope. For confocal imaging, the laser powers and corresponding average intensities were set as follows: Panels A, E, I and M, the 561 nm laser power was set at 15 μW (~ 0.3 W/cm^2^) and the 405 nm laser intensity ranged from ~ 0.2 - 0.8 W/cm^2^; Panels B, F, J, and N, the 405 nm laser power was set at 22 μW (~ 0.2 W/cm^2^) and the 561 nm laser intensity ranged from ~ 0.15 - 0.5 W/cm^2^. Four FOVs were taken for each condition with cells expressing mEos3.2. Two FOVs were taken for each condition with wildtype cells. For widefield imaging, 561 nm laser intensity was set at 1 kW/cm^2^ for panels C, G, K, O; 405 nm laser intensity was set at 1 W/cm^2^ for panels D, H, L, P. Eight to ten FOVs were taken for each condition with cells expressing mEos3.2. Three or four FOVs were taken for each condition with wildtype cells. Eq. 7 of the 3-state model was fit to the time courses of fluorescence signal to determine the parameters giving the best fit: **(A-D)** The product of total number of molecules per cell and the signal of an R-state mEos3.2 molecule per frame (M_n_ x ε_f_); **(E-H)** photoconversion rate constant (k_act_); and **(I-L)** photobleaching rate constant (k_bl_) (Table S3, S4). **(M-P)** Time-integrated signal per cell 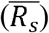 calculated using Eq. 8 (Table S3, S4). The error bars in the left 2 columns are 95% confidence intervals of the fit. The error bars in the right 2 columns are the weighted standard deviations among different FOVs. Fig. S8 and S9 report the raw data.

To compare imaging conditions quantitatively, we explored the effects of laser intensities on mEos3.2 photophysics by point-scanning illumination (Fig. 4, left 2 columns, Table S3). 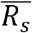 decreased (Fig. 4M), and both rate constants (Fig. 4E, I) increased with higher 405-nm laser intensity. 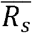 increased with higher 561-nm laser intensity (Fig. 4N), but the 561-nm laser intensity had only modest effects on both rate constants (Fig. 4F, J) in the range we tested.

We used widefield illumination to explore the effects of a wider range of 561-nm laser intensities on mEos3.2 photophysics, including the high 561-nm laser intensity of ~1 kW/cm^2^ often used in SMLM (Fig. 4, right 2 columns, Table S4). Values of k_bl_ increased with higher 405-nm laser intensity (Fig. 4K), as observed with point-scanning illumination (Fig. 4I), but the 405-nm laser intensity had remarkably little impact on k_act_ (Fig. 4G) under SMLM conditions, which was likely caused by the high 561-nm laser intensity of 1 kW/cm^2^ driving some photoconversion. The time-integrated signal per cell 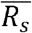 increased with higher 561-nm laser intensity and then plateaued and dropped (Fig. 4P). Both k_act_ and k_bl_ increased dramatically with 561-nm laser intensities (Fig. 4H, L) above the intensities used for point-scanning illumination (Fig. 4F, J). The discussion section considers these differences between the confocal and widefield conditions.

### Photon counts from single red-state mEos3.2 molecules in live and fixed *S. pombe* cells

To assess the effects of fixation and the imaging buffer on mEos3.2 photophysics under SMLM conditions, we compared the photon counts of single R-state mEos3.2 molecules per frame in live and fixed yeast cells (Fig. 5) under SMLM imaging conditions with a frame rate of 50 fps and a 561-nm laser intensity of 1 kW/cm^2^. We localized the single R-state molecules in each frame and measured their photon counts when the density of the R-state molecules was sparse enough for localization.

**Figure 5:**
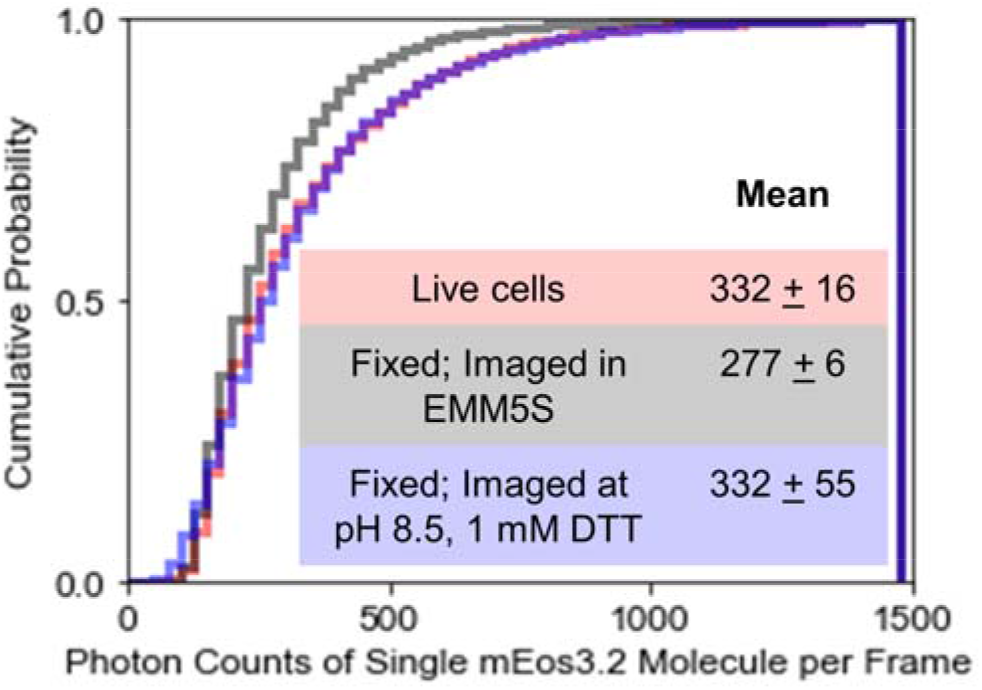
The effect of fixation and imaging buffer on photon counts from mEos3.2 by single molecule localization microscopy (SMLM). SMLM imaging of cytoplasmic mEos3.2 in *S. pombe* cells with continuous illumination at 405 nm (1 W/cm^2^) and 561 nm (1 kW/cm^2^) under 3 conditions: live cells (red), cells fixed with 2% formaldehyde in EMM5S medium for 30 min and mounted in EMM5S synthetic medium (black) or mounted in 50 mM Tris-HCl (pH 8.5) with 1 mM DTT) (blue). Four FOVs of 40 μm x 40 μm were taken over time at 50 fps for 15,000 frames for each condition. All emission bursts between frame 5,000 and 10,000 were localized to measure the photon counts from single red mEos3.2 molecules in each 20-ms frame, when the mEos3.2 molecules were sparse enough for Gaussian center fitting. The curves show the cumulative probability distribution of the photon counts of single mEos3.2 molecules in each 20-ms frame under all three conditions. Inset: The table reports the mean number of photons (+ standard deviation between the 4 FOVs) emitted by single mEos3.2 molecules in each 20-ms frame under the three conditions. About 2 – 5 x 10^5^ emission bursts were recorded for the histogram.

The mean photon counts per frame from single R-state mEos3.2 molecules in live cells and fixed cells imaged in the Tris8.5-DTT buffer were identical (P = 1.000) and higher than the counts in fixed cells imaged in the EMM5S medium at pH 5.5 (P = 0.007, Fig.5, inset). Therefore, the higher brightness per frame of R-state mEos3.2 molecule in the fixed cells in Tris8.5-DTT buffer can improve localization precision (35). Besides, the oxidizing environment in fixed cells in EMM5S medium may prolong the total on-time (the total time a mEos3.2 molecule is in the red fluorescent state until bleached) of single R-state mEos3.2 molecules, suggested by the time-integrated signal per cell 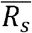 (the product of brightness per frame and total on-time of single R-state mEos3.2, and the total number of molecules per cell) being higher in the fixed cells in EMM5S medium than in the other 2 conditions (Fig. 4O).

### Intermediate state of mEos3.2 that converts to the red fluorescent state upon 561-nm illumination

To investigate the effects of 405-nm and 561-nm illumination separately, we alternated periods of multiple cycles of 405-nm and 561-nm illumination. Each period of 405-nm illumination increased the fluorescence emission during 561-nm illumination; surprisingly, the signal rose transiently above the initial value despite the 405-nm illumination being switched off (Fig. 6A). After peaking during the fifth cycle of each 561 nm illumination period, the signal decreased due to photobleaching (Fig. 6A, B). The signal during each 561-nm illumination period (except for the first one without prior 405-nm illumination) followed similar time courses (Fig. 6B). These transient increases in the fluorescence signal also occurred using alternating widefield illumination conditions (Fig. S4).

**Figure 6:**
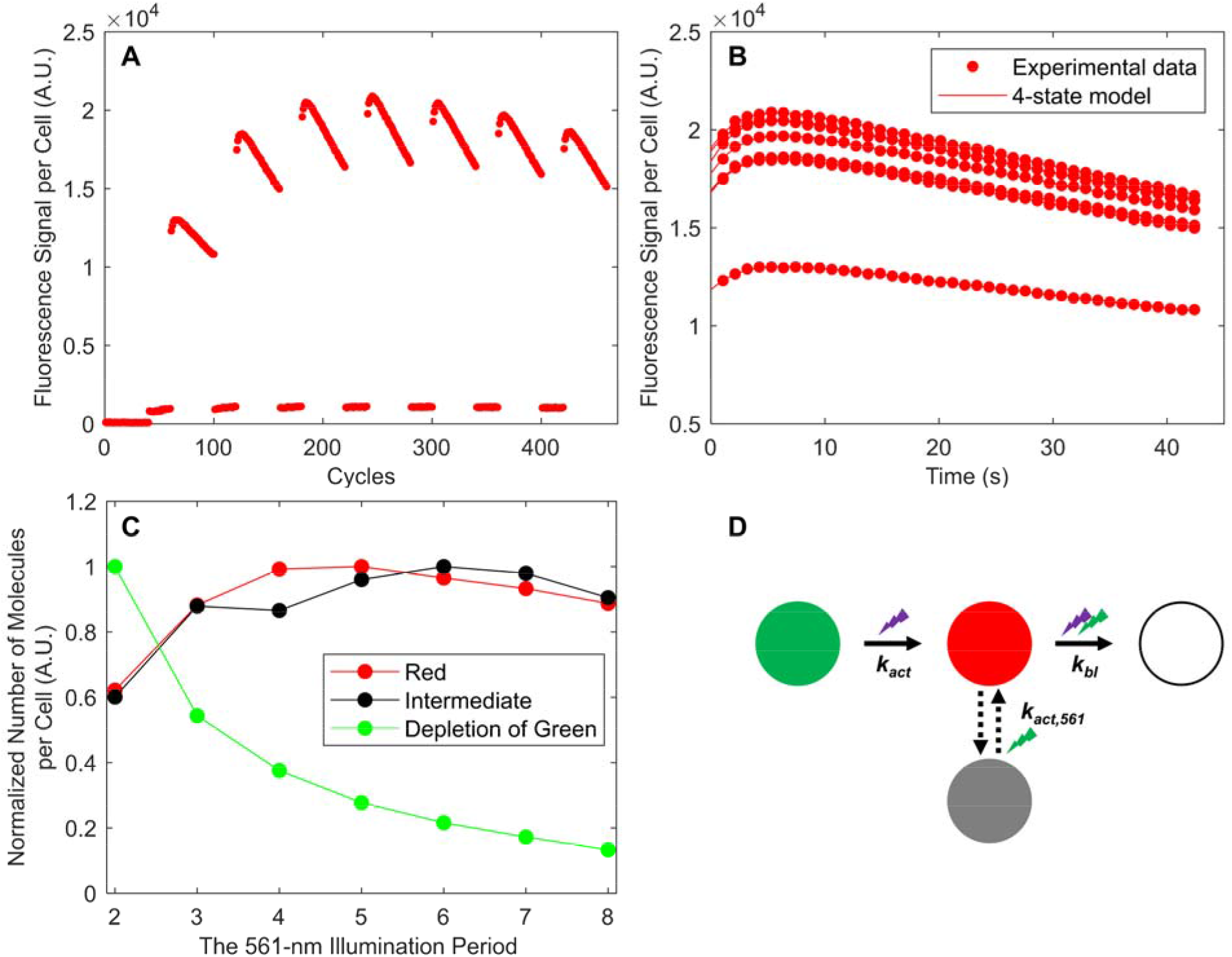
An intermediate state of mEos3.2 that requires 561-nm illumination to reach the red-state. **(A)** Time courses of the fluorescence signal per live *S. pombe* cell expressing mEos3.2 and subjected to alternating illumination by point-scanning confocal microscopy at 561 nm (37 μW) for 40 cycles followed by illumination at 405 nm (56 μW) for 20 cycles. Illumination at 561 nm increased the fluorescence signal beyond the start of the 561-nm illumination period after each period of 405-nm illumination. **(B)** Comparisons of the time courses of the fluorescence signals during 7 periods with 40 cycles of 561-nm illumination (not including the first 561-nm illumination period before any 405-nm irradiation) from panel A. Eq. 14 of the 4-state model in panel D (lines) was fit to these time courses (dots) to determine rate constants giving the best fit. The mean activation rate constant (k_act,561_) from the I-state to the R-state is 0.34 s^-1^ (SD: 0.02).. **(C)** Normalized number of red (red), intermediate (black) and green molecules (green) at the beginning of each 561-nm illumination period. The normalized number of red and intermediate molecules were estimated from fitting the Eq. 14 of the 4-state model to the time courses of fluorescence signal during the 7 periods of 561-nm illumination in Panel B. The depletion of green molecules was estimated from subtracting the fluorescence signal at the last cycle of the previous 561-nm illumination period from the signal at the first cycle of the previous 561-nm illumination period. **(D)** Four-state model for mEos3.2 photoconversion and bleaching. Illumination at 405 nm photoconverts mature mEos3.2 molecules from the G-state to the R-state with a photoconversion rate constant of k_act_. A subpopulation of the R-state molecules enter an intermediate state (gray circle), and irradiation at 561 nm activates the I-state mEos3.2 molecules to the R-state with a rate constant of k_act,561_. Illumination at both wavelengths photobleaches red mEos3.2 molecules with a rate constant of k_bl_. See Fig. S4 and S5 for supplementary information.

We considered four hypotheses to explain these transient increases in the fluorescence signal. First, illumination at 561 nm might photoconvert G-state mEos3.2 to the R-state directly, but we observed no comparable activation with 561-nm irradiation alone (Fig. S5B, and first 40 cycles of 561-nm irradiation in Fig. 6A). A second hypothesis is that the first-order photoconversion reaction is slow after absorption of 405-nm photons, delaying accumulation of R-state molecules. We ruled out this mechanism by adding a 2-min pause between each 405-nm illumination period and the following 561-nm illumination period. The transient increase in the fluorescence signal was still observed, ruling out this hypothesis as a dominant effect (Fig. S5A). Similarly, we could rule out the third hypothesis that the observed increase in the fluorescence signal over the course of 561-nm illumination was related to protein maturation or similar live-cell phenomena by observing the same effect in fixed cells (Fig. S5C).

The fourth hypothesis is that 405-nm illumination leaves a subpopulation of mEos3.2 molecules in an intermediate (I) state that requires 561-nm illumination to convert to the fluorescent R-state. To test this hypothesis, we added a fourth I-state to the model (Fig. 6D). Whether this I-state is populated from the R-state or G-state mEos3.2 molecules does not affect Eqs. 9–14, which only consider the period of 561-nm illumination when conversion into the I-state is negligible. Eq. 14 of our 4-state model fits closely the time courses of mEos3.2 fluorescence signal during each 561-nm illumination period (Fig. 6B). The best fits gave an average activation rate constant for conversion from the I-state to R-state by 561-nm irradiation k_act,561_ = 0.34 s^-1^ (SD: 0.02 s^-1^) (Fig. 6B), which is 4-fold higher than the rate constant for photoconversion from the G- to R-state, k_act_ = 0.050 s^-1^ (95% CI: 0.048-0.051 s^-1^), as measured with simultaneous 405-nm and 561-nm illumination (Fig. S5D).

We compared the number of I-state molecules changing over time with the respective G- and R-state populations (Fig. 6C). The normalized time course of the I-state molecules was similar to R-state molecules and distinctly different from the G-state molecules, strongly suggesting that the I-state is populated from the R-state molecules. Overall, these experiments revealed that a subpopulation of the R-state molecules is converted to the I-state but can be converted back to the R-state by 561-nm illumination (Fig. 6D). We did not detect any spontaneous conversion from I- to R-state over 2 minutes (Fig. S5A). However, for experiments with simultaneous illumination at both 405 nm and 561 nm, Eq. 7 of our simplified 3-state model (without considering the I-state) fully accounted for the time courses of fluorescence signal per cell under all the conditions (Fig. 1B, 2C, 3A, S8, S9).

## DISCUSSION

### Characterizing the photophysics of mEos3.2 by fitting our 3-state model to quantitative time-lapse fluorescence microscopy data

We show how to measure the product of total number of molecules per cell and the signal per R-state molecule per frame (M_n_ x ε_f_) and rate constants for photoconversion (k_act_) and photobleaching (k_bl_) in cells by fitting a simple 3-state model to the time course of measured fluorescence signals in the red channel (Fig. 1). Eq. 7 of our 3-state model fit the time courses of the fluorescence signal per cell very closely (Fig. 2C, 3A) and revealed how different conditions changed the three parameters separately. Furthermore, simulations of the 3-state model showed how the values of the three parameters influenced the time courses of the number of R-state molecules per cell (Fig. S2B – D). We also used Eq. 8 to calculate the time-integrated signal per cell 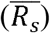 using M_n_ x ε_f_ and k_bl_.

Our approach has several advantages over the alternative approach of characterizing single fluorescent molecules *in vitro* (18, 37, 38). (1) It avoids potential artifacts caused by using arbitrary photon number or localization uncertainty thresholds in single-molecule localization algorithms to separate molecules from noise. (2) It is easy to implement with conventional microscopes and whole cells. (3) Large numbers of cells can be imaged in just hours to test different sample preparation and imaging conditions, including a wide range of laser intensities. (4) It extracts photoconversion or photoactivation rate constants from PAFPs or PCFPs in cells more easily than single-molecule methods, as photobleaching is hard to account for in single-molecule data (30). These rate constants are useful for optimizing SMLM imaging conditions and simulating SMLM raw data.

On the other hand, our approach cannot measure the single-molecule blinking kinetics of mEos variants in either their green (39, 40) or red states (41, 42). Blinking in the green state of mEos variants could affect photoconversion to the red state (39, 40). The interplay between blinking and photoconversion has also been described in other photoconvertible fluorescent proteins, such as SAASoti (43) and LEA (44). Thus, combining our approach with single-molecule measurements will offer a more complete and quantitative understanding of the photophysics of PAFPs or PCFPs.

### Sample preparation for imaging mEos3.2 in fixed fission yeast cells

Preserving the fluorescence signal and structures of interest is crucial when fixing cells with fluorescent protein labels. We show that formaldehyde fixation does not destroy mEos3.2 molecules but permeabilizes yeast cells for small molecules, making the photophysical properties of mEos3.2 sensitive to the composition of the imaging buffer. The low pH of 5.5 and lack of oxygen scavenging system in the EMM5S synthetic medium affected the photophysical properties of mEos3.2 in fixed cells as expected from previous works showing that EosFP photoconverts faster (4) and mEos3.2 emits fewer red photons at acidic pH (6). Photooxidation can increase the photobleaching rate (45). Oxygen in the solution can also affect the fluorescence signal by promoting intersystem crossing (46) and convert the excited molecules to the non-fluorescent triplet state.

Our experiments also show that fixation conditions should be tested and optimized for each fluorescent protein. For example, mEos3.2 had even lower 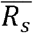 and higher k_act_ and k_bl_ in cells fixed for 30 min rather than 15 min under the tested imaging condition (Fig. 2E-G). On the other hand, GFP was far less sensitive to fixation, as formaldehyde treatment had little effect on its fluorescence signal and photobleaching rate (Fig. S6).

The photophysics of mEos3.2 was similar in live cells and fixed cells in the Tris8.5-DTT buffer (Fig. 3). Imaging buffers with a pH equal to or slightly higher than the cytoplasmic pH of fission yeast at ~7.3 (47) increased 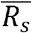 and reduced k_act_ and k_bl_ (Fig. 3B, D, E). Adding a reducing agent DTT to the imaging buffer further increased 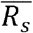 (Fig. 3B). The Tris8.5-DTT buffer not only maintained the photon counts of single R-state mEos3.2 molecules in each frame (Fig. 5) but also reduced the cellular autofluorescence compared with fixed cells in EMM5S medium (Fig. S8E, S8H). This is crucial for SMLM imaging, where the signal-to-noise ratio is important for obtaining high localization precision and consequently resolution (37).

### Comparison of point-scanning and widefield illumination for mEos3.2 photophysics

We compared the photophysical properties of mEos3.2 under point-scanning and widefield illumination. Under point-scanning illumination, each area of the sample was illuminated for a very short time at high peak intensity (e.g. ~80 kW/cm^2^, Fig. 4), while the other pixels were kept in the dark. Thus, the average intensity of the laser power over the entire field of view was ~10^4^ times lower than the peak intensity (e.g. ~0.5 W/cm^2^, Fig. 4).

Despite huge differences in the instantaneous peak intensities in point-scanning and widefield microscopy, the rate constants for photoconversion and photobleaching in live cells were similar at comparable average intensities (Table S3, S4). For example, with average intensities of ~ 0.5 to 1.1 W/cm^2^ for both lasers, confocal and widefield imaging of live cells gave similar values for k_act_ (~1.5 x 10^-2^ s^-1^ vs. ~5 x 10^-2^ s^-1^) and k_bl_ (~2 x 10^-3^ s^-1^ vs. ~1 x 10^-3^ s^-1^). Moreover, the photophysical parameters of mEos3.2 trended similarly with illumination intensities by both point-scanning confocal and widefield microscopy. For example, k_bl_ increased with higher 405-nm intensity (Fig. 4I, K) and M_n_ x ε_f_ increased with higher 561-nm laser intensity (Fig. 4B, D). However, these trends diverged at higher 561-nm laser intensities of 100 and 1000 W/cm^2^ above the range of the confocal microscope. For example, k_act_ increased dramatically with 561-nm laser intensity (Fig. 4H), but only increased slightly with 405-nm laser intensity (Fig. 4G, 561-nm laser intensity = 1 kW/cm^2^). We conclude that the 561-nm laser contributed strongly to photoconversion at high intensities.

The effects of the fixation and imaging buffer on mEos3.2 photophysics is also consistent under different illumination methods. For example, under comparable, low average 405-nm and 561-nm laser intensities (~1 W/cm^2^), the time-integrated signal per cell 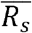 (Fig. 4N, P) was lower and photoconversion (Fig. 4F, H) and photobleaching rates constants (Fig. 4J, L) were higher in fixed cells in EMM5S than in live cells and fixed cells in the Tris8.5-DTT buffer. However, fixation and imaging buffer affect mEos3.2 photophysics differently at higher 561-nm laser intensities of 100 and 1000 W/cm^2^, above the range of confocal microscope. For example, 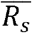 was lower in fixed cells in EMM5S (pH 5.5) than in live cells under low 561-nm laser intensity in the range of 1-10 W/cm^2^, while for a high 561-nm laser intensity of 1 kW/cm^2^ the trend was the opposite (Fig. 4P). The 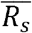 in the fixed cells was only higher in the Tris8.5-DTT buffer than in EMM5S under low 561-nm laser intensities of 1-10 W/cm^2^ (Fig. 4P). Moreover, under low 561-nm laser intensities both k_act_ (Fig. 4E, H) and k_bl_ (Fig. 4I, L) in fixed cells in EMM5S were higher than in live cells and fixed cells in Tris8.5-DTT buffer, while under higher 561-nm laser intensities some measurements of the rates were lower than in the Tris8.5-DTT buffer (Fig. 4H, L). Oxygen in the environment could promote intersystem crossing and convert the excited fluorophores to the non-fluorescent triplet state, where the molecules could return to the fluorescent state through laser excitation. The laser intensity could potentially affect the probability that the triple state molecules return to the fluorescent state. Therefore, redox environment in the imaging buffer for fixed cells may affect mEos3.2 photophysics differently depending on the 561-nm laser intensity.

### Comparing laser intensities for imaging mEos3.2 in live and fixed fission yeast cells

Our quantitative measurements provide information for selecting laser intensities to image proteins tagged with mEos3.2. Maximizing the red fluorescence signal of mEos3.2 while maintaining a relatively low level of autofluorescence background is the key to optimize imaging quality. Higher signal-noise-ratios can increase localization precision and thus the resolution in SMLM (35). For SMLM, one controls the rates of photoconversion and photobleaching to achieve a density of active fluorophores sparse enough for localization but also dense enough to image quickly. High laser intensities are usually used for fast SMLM imaging in fixed samples (37).

Higher 405-nm illumination intensity had four effects relevant for SMLM image quality: (1) it decreased 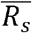 under a low average 561-nm laser intensity of ~ 0.3 W/cm^2^ (Fig. 4M), but the effect was not obvious under high 561-nm laser intensity of 1 kW/cm^2^ (Fig. 4O); (2) it increased background autofluorescence (Fig. S8B, E, H), especially in live cells (Fig. S8B); (3) it increased k_act_ (Fig. 4E); and (4) it increased k_bl_ (Fig. 4I, K). One may ramp up the 405-nm laser intensity while imaging a FOV to increase k_act_ and compensate for the loss of bleached molecules. However, high 405-nm laser intensities can potentially decrease SMLM imaging quality in two ways: increased autofluorescence compromises accurate localization; and rapid photobleaching decreases 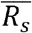, likely decreasing the total number of localizations.

The time-integrated signal 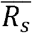 increased over the low range of 561-nm laser intensities (Fig. 4N) and peaked in the middle of the high range of 561-nm laser intensities (Fig. 4P), so very high 561-nm laser intensities could compromise image quality. Moreover, both the photoconversion (Fig. 4H) and photobleaching rates (Fig. 4L) increased at high 561-nm laser intensities. The photoconversion rate likely increases, because 561-nm laser is the predominant source of photoconversion at high intensities (Fig. 6D). Our results differ from Thedie *et al*. (39), who reported that high 561-nm laser intensities (1.2 – 4.8 kW/cm^2^) slow the photoconversion of single mEos2 molecules embedded in PVA under weak 405-nm illumination (0.03 W/cm^2^). Their interpretation was that 561-nm illumination pushes green-state mEos2 molecules into a transient off state that cannot be photoconverted. Both the sample conditions and illumination intensities differ in the 2 experiments. Their mEos2 molecules were immobilized in PVA with restricted access to oxygen rather than being in a physiological environment in cells. Additionally, we used higher 405-nm illumination (1 W/cm^2^) and lower 561-nm laser intensity (0.001 – 1 kW/cm^2^) where green molecules in the transient off state may convert back to the fluorescent state, thus promoting photoconversion to the red-state.

Based on our data, we recommend the following for SMLM imaging of mEos3.2 in live and fixed fission yeast cells. (1) Use high but not maximal 561-nm laser intensities to avoid decreasing 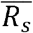 (Fig. 4P) and degrading the SMLM image quality. (2) Use low 405-nm laser intensities to achieve sparse activation for temporal separation of molecules and to compensate for the high photoconversion rate from strong 561-nm laser illumination.

### Application of our findings to quantitative SMLM with mEos3.2 in live and fixed yeast cells

Several variants of EosFP fluorescent proteins are used for SMLM of live and fixed cells (9, 11, 12). Both sample preparation and imaging conditions can affect quantitative measurement. The time-integrated signal 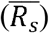 contains information of both the brightness per single fluorescent molecule and the total number of molecules able to fluoresce in the structure of interest. The single-molecule brightness also affects the number of mEos3.2 localizations, as only molecules emitting more than a threshold number of photons in each frame are counted (34). Under SMLM imaging conditions (Fig. 4O), 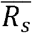 measurements differed in live cells and fixed cells in the Tris8.5-DTT buffer, so they cannot be used for direct comparisons. However, the photon counts from single mEos3.2 emission bursts in each SMLM frame were similar in live cells and fixed cells (Fig. 5), so the same analysis thresholds can be used for single-molecule localizations. Combining the time-integrated signal 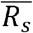 (Fig. 4O) and photon counts (Fig. 5, inset) suggests that fixation and the imaging buffer affect the total on-time of single R-state mEos3.2, likely due to the oxidation-reduction environment. Similarly, adding reducing thiol MEA and removing oxygen from the imaging buffer increase the number of blinking cycles of R-state mEos2 in fixed mammalian cells (41) and of purified mEos3.2 (48).

### Illumination at 561-nm converts intermediate-state mEos3.2 to the red fluorescent state

Our alternating illumination experiments (Fig. 6) revealed that some R-state molecules enter the I-state but can be reconverted back to the R-state by 561-nm illumination. De Zitter *et al*. has reported a similar observation, that mEos4b – an mEos3.2 derivative – recovered from a long-lived dark state to the red state in response to 561-nm illumination (42). The long-lived dark state recovery rate increased with 561-nm laser intensity and could be recovered to the red state by 405-nm irradiation. Our I-state converts to R-state upon 561-nm illumination, even at as low as 1 W/cm^2^ (Fig. S4A). However, our I-state molecules did not seem to respond to 405-nm illumination. The period of 561-nm illumination would not increase the fluorescence signal transiently (Fig. 6A), if 405-nm irradiation could convert the I-state to R-state. Therefore, the I-state we observed for mEos3.2 is unlikely to be the long-lived dark state as described for mEos4b (42). Further structural studies are needed to characterize the nature of this intermediate state.

## CONCLUSION

We measured the time-integrated signal per cell and the rate constants for photoconverting and photobleaching of mEos3.2 in fission yeast cells by fitting a 3-state model to time-lapse quantitative fluorescence microscopy data. Formaldehyde fixation partially permeabilized the yeast cell membrane for small molecules, allowing the reducing agent and ions in the imaging buffer to enter fixed yeast cells and affect mEos3.2 photophysics. Using an imaging buffer at pH 8.5 with 1 mM DTT resulted in a time-integrated signal comparable to live cells under certain imaging condition, suggesting that mEos3.2 molecules survived the light chemical fixation. We investigated the effects of fixation and imaging buffer on mEos3.2 photophysics in yeast cells over a wide range of laser intensities by point-scanning and widefield microscopy. We discovered an intermediate state of mEos3.2, populated by red-state molecules, that can be convert to the red fluorescent state by 561-nm illumination. Our results provide information for sample preparation and laser intensities for imaging and counting mEos3.2-tagged molecules in fission yeast cells. Our imaging assay and 3-state model can also be applied to study the photophysical properties of other photoactivatable or photoconvertible fluorescent proteins.

## SUPPORTING MATERIAL

Supporting Material can be found online at ***.

## AUTHOR CONTRIBUTIONS

HS, JB and TDP designed research; HS carried out the experiments; KH built the single-molecule localization microscope and assisted in imaging and data analysis; all authors analyzed the data and wrote the paper.

## ACKNOWLEDGMENTS

Research reported in this publication was supported by National Institute of General Medical Sciences of the National Institutes of Health under award number R01GM026132 and R01 GM118486, and the Wellcome Trust (203285/B/16/Z). MS and KH were supported by NIH Grant No. T32EB019941. The content is solely the responsibility of the authors and does not necessarily represent the official views of the National Institutes of Health. The authors thank Bennett Rollins and Zach Marin for help with single-molecule localization imaging and data analysis, Yongdeng Zhang for helpful discussion, and Phylicia Kidd for help with sample preparation. We also acknowledge support from the Program in Physics, Engineering and Biology. JB discloses a significant financial interest in Bruker Corp. and Hamamatsu Photonics.

