## Supplementary Information for "Sample preparation and imaging conditions affect mEos3.2 photophysics in fission yeast cells"

**Supplementary Figure 1**

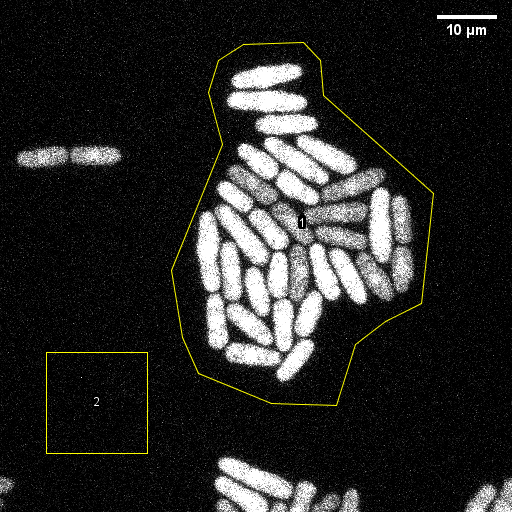

**Supplementary Figure 1: Measurement of mEos3.2 fluorescence signal per cell by quantitative fluorescence microscopy and analysis with Fiji.** Fluorescence micrograph of a field of *S. pombe* cells expressing cytoplasmic mEos3.2 at the 31^st^ time cycles as in Fig. 1. Region of Interest (ROI) 1 containing cells was manually selected with a polygon tool, and ROI 2 was manually selected with a square tool for background subtraction. Scale bar = 10 µm.

**Supplementary Figure 2**

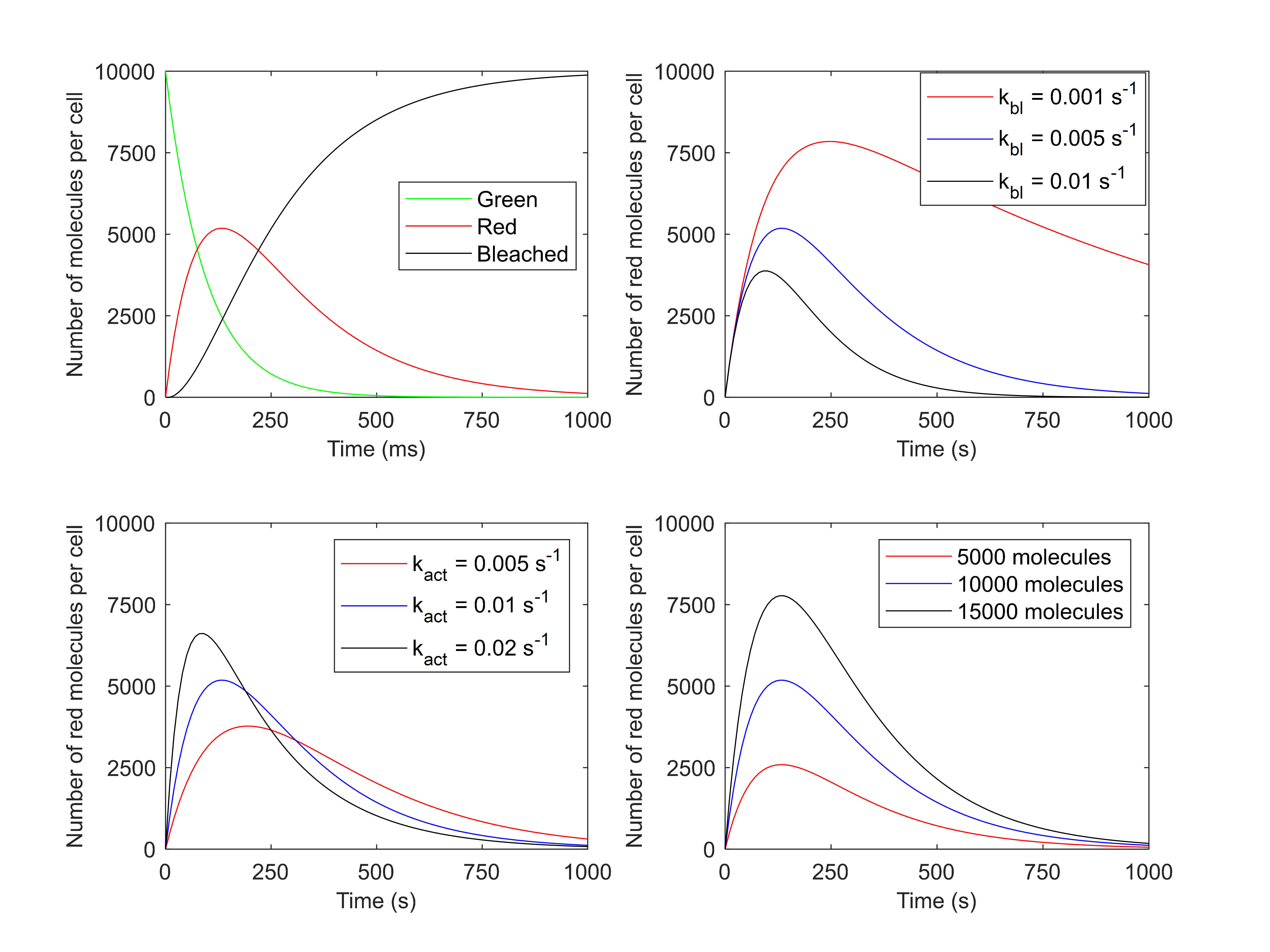

**Supplementary Figure 2: Simulations of the numbers of green, red, and bleached mEos3.2 molecules per cell over time. (A)** Evolution of the number of the green, red, and bleached mEos3.2 molecules. Conditions: 10,000 total molecules per cell, photoconversion rate constant k_act_ = 0.01 s^-1^, photobleaching rate constant k_bl_ = 0.005 s^-1^. **(B)** Vary the photobleaching rate constant. Conditions: 10,000 total molecules per cell, k_act_ = 0.01 s^-1^. **(C)** Vary the photoconversion rate constant. Conditions: 10,000 total molecules, k_bl_ = 0.005 s^-1^. **(D)** Vary the total number of molecules per cell. Conditions: k_act_ = 0.01 s^-1^, k_bl_ = 0.005 s^-1^.

**Supplementary Figure 3**

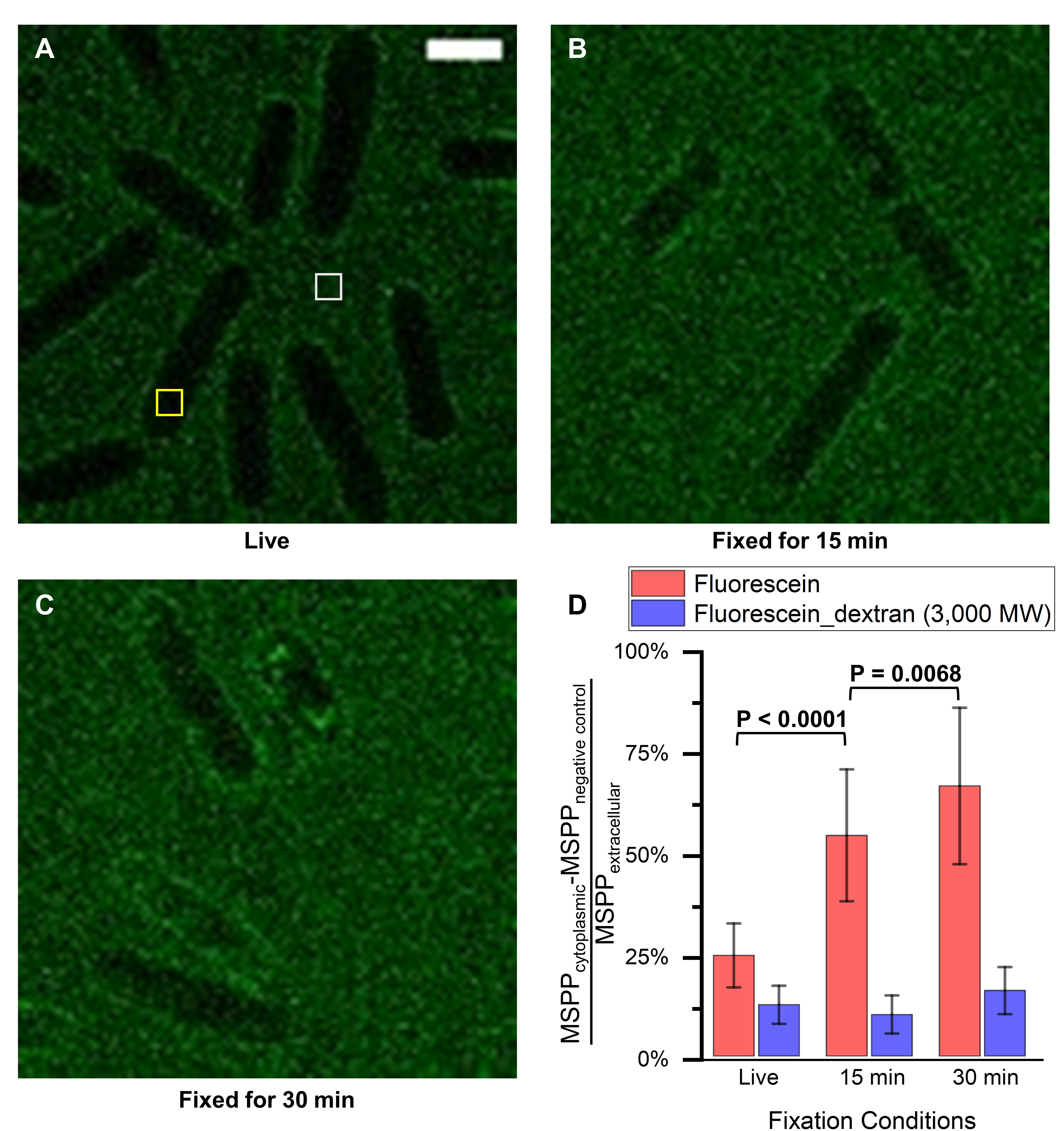

**Supplementary Figure 3: Effect of formaldehyde fixation on the permeability of *S. pombe* cells. (A-E)** Fluorescein mid-section images (green) of wild type *S. pombe* cells incubated in 20 µM fluorescein diluted in EMM5S medium. Conditions: **(A)** Live cell (scale bar = 5 µm, yellow box: ROI for measuring cytoplasmic mean signal per pixel (MSPP), white box: ROI for measuring extracellular mean signal per pixel); **(B)** cells fixed with 2% formaldehyde for 15 minutes; **(C)** cells fixed with 2% formaldehyde for 30 minutes. **(D)** Quantify cell permeability in live and fixed wildtype *S. pombe* cells using fluorescein and fluorescein_dextran (3,000 MW). Average cytoplasmic mean signal per pixel (MSPP_cytoplasmic_) after autofluorescence subtraction (MSPP_negative control_) divided by extracellular mean signal per pixel (MSPP_extracellular_) were plotted. The error bars were standard deviation between cells in each condition. There was a significant percentage increase in the fluorescein signal in the cells fixed with 2% formaldehyde for 15 min compared to the live cells (P<0.0001). Longer fixation time (30 minutes versus 15 minutes, P = 0.0068) also increased the fluorescein signal in cells.

**Supplementary Figure 4**

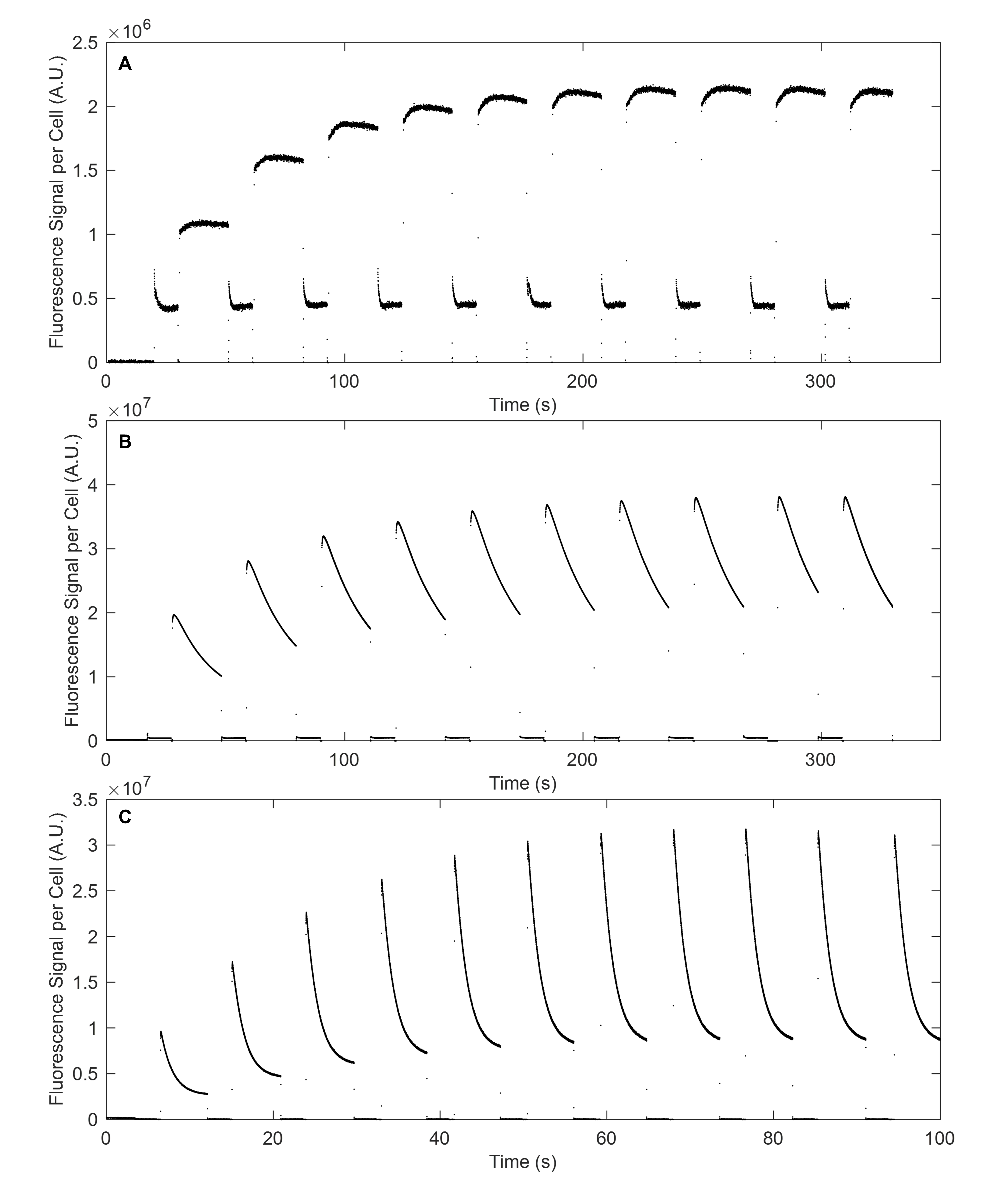

**Supplementary Figure 4: The intermediate state that converts to the red fluorescent state under wide-field illumination at 561 nm.** Time courses of the fluorescence signal per live *S. pombe* cell expressing mEos3.2 and subjected to alternating widefield illumination by at 561 nm followed by illumination at 405 nm at 1 W/cm^2^. Illumination at 561 nm increased the fluorescence signal beyond the start of the 561-nm illumination period after each period of 405-nm illumination. **Conditions:** **(A)** 561 nm laser intensity = 1 W/cm^2^, 50 fps; 20 s for each 561-nm illumination period, followed by 10 s for each 405-nm illumination period **(B)** 561 nm laser intensity = 10 W/cm^2^, 50 fps; 20 s for each 561-nm illumination period, followed by 10 s for each 405- nm illumination period **(C)** 561 nm laser intensity = 100 W/cm^2^, 500 fps; 5 s for each 561-nm illumination period, followed by 2.5 s for each 405- nm illumination period.

**Supplementary Figure 5**

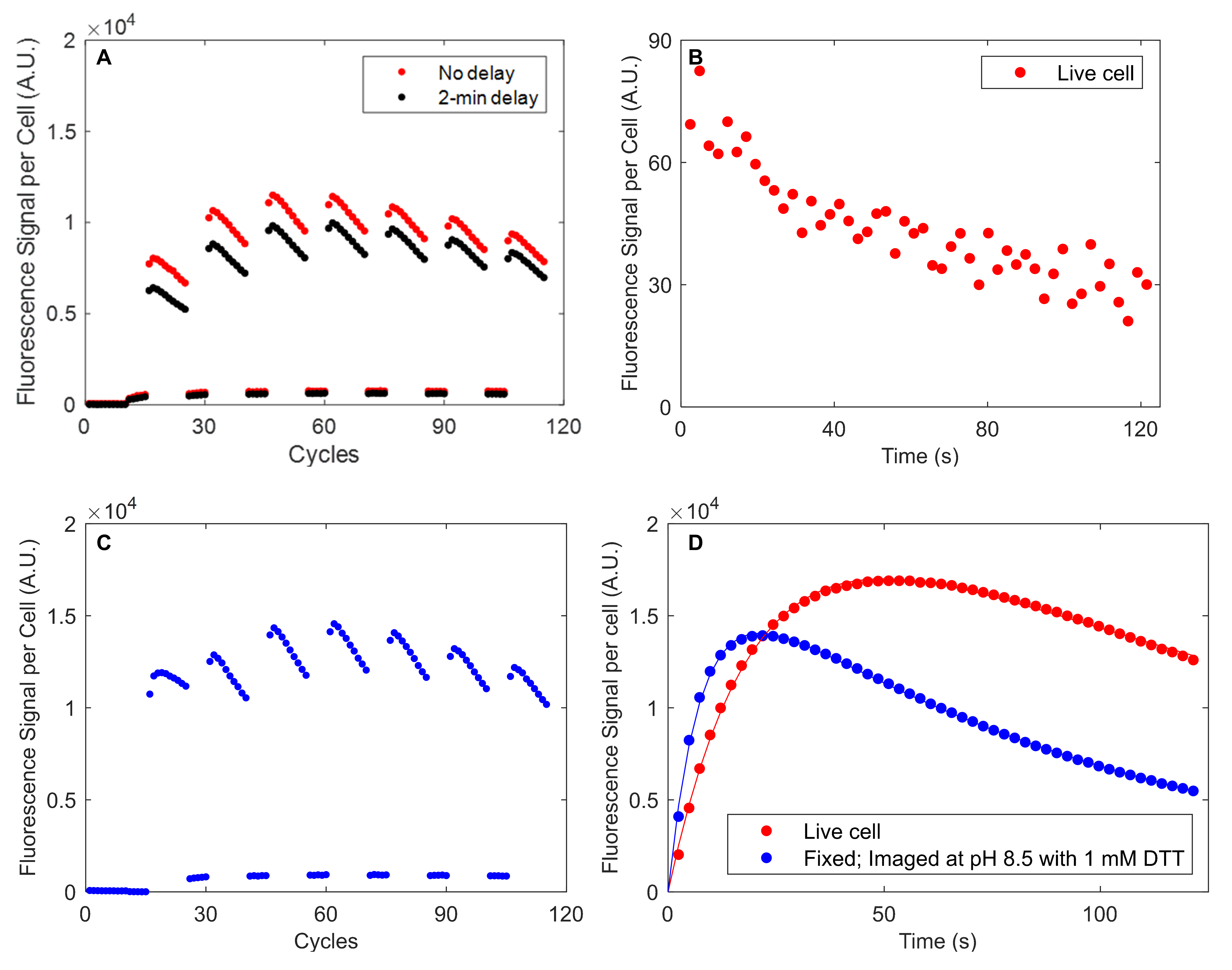

**Supplementary Figure 5:** **The intermediate state that converts to the red fluorescent state by 561-nm illumination. (A)** Time courses of the fluorescence signal per live *S. pombe* cell expressing mEos3.2 and subjected to alternating illumination by point-scanning confocal microscopy at 561 nm (37 µW) for 10 cycles followed by illumination at 405 nm (56 µW) for 5 cycles. Illumination at 561 nm increased the fluorescence signal beyond the start of the 561-nm illumination period after each period of 405-nm illumination. The red dots are data collected from live cells with no delay between the periods of illumination at 405 nm and 561 nm, while the black dots are data collected from live cells with a 2-min break after each period of 405-nm illumination before the following period of 561-nm illumination. Fitting Eq. 14 of the 4-state model in panel D fit to these time courses of the fluorescence signals during 7 periods with 10 cycles of 561-nm illumination gave the mean activation rate constant (k_act,561_) from the I-state to the R-state of 0.27 s^-1^ (SD: 0.03) for the experiment with no delay between the 405-nm and 561-nm illumination cycle (red), and of 0.28 s^-1^ (SD: 0.03) for the experiment with 2-min breaks (black). **(B)** Time course of the fluorescence signal per cell expressing mEos3.2 and illuminated only at 561 nm (37 µW). **(C)** Time course of the fluorescence signal per cell expressing mEos3.2, fixed with 2% formaldehyde for 30 min, and imaged in the buffer at pH 8.5 with 1 mM DTT under alternating illumination by point-scanning confocal microscopy at 561 nm (37 µW) for 10 cycles followed by illumination at 405 nm (56 µW) for 5 cycles. The mean activation rate (k_act,561_) of mEos3.2 from the I-state to the R-state is 0.27 s^-1^ (SD: 0.03). **(D)** Time courses of the fluorescence signal per cell from live *S. pombe* cells and cells fixed with 2% formaldehyde in EMM5S medium for 30 min and imaged in 50 mM Tris-HCl buffer at pH 8.5 with 1 mM DTT, and illuminated simultaneously at 405 nm (56 µW) and 561 nm (37 µW). Eq. 7 of the 3-state model (line) was fit to the time courses of the fluorescence signal per cell (red or blue dots). The rate constants of photoconversion from the G-state to the R-state (k_act_) were 0.050 s^-1^ (95% CI: 0.048-0.051) for live cells (red) and 0.131 s^-1^ (95% CI: 0.127-0.134) for fixed cells (blue).

**Supplementary Figure 6**

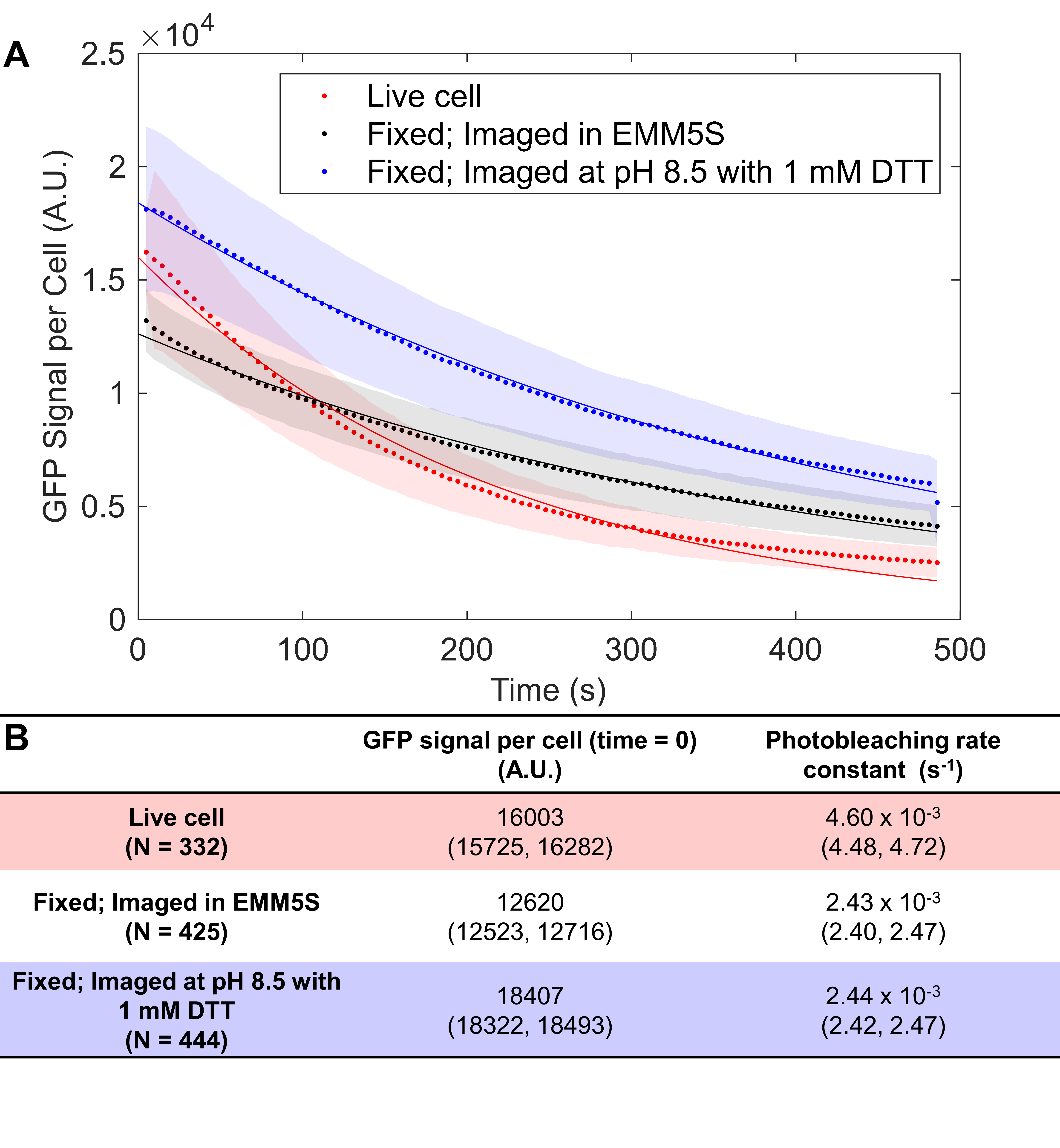

**Supplementary Figure 6: Effects of fixation and imaging buffer on the fluorescence signal and photobleaching of GFP in *S. pombe* cells. (A**) Time courses of the GFP fluorescence signal per *S. pombe* cell expressing Fim1-GFP and excited at 488 nm by point-scanning confocal microscopy under 3 conditions: (red dots) live cells; (black dots) fixed with 2% formaldehyde in EMM5S for 30 min and imaged in EMM5S; (blue dots) fixed with 2% formaldehyde in EMM5S for 30 min and imaged in 50 mM Tris-HCl buffer at pH 8.5 with 1 mM DTT. Z-stacks from 4 FOVs of 85 x 85 µm were recorded for Fim1-GFP cells and 2 FOVs were recorded of wild type cells for autofluorescence background subtraction. A single exponential decay was fit to the time course of fluorescence signal (dots). The lines are theoretical curves with rate constants giving the best fit to the data. **(B)** Values of the GFP signal per cell at time 0 (without photobleaching) and the photobleaching rate constant from the best fit to the experimental data. The 95% confidence intervals of the fitted parameters are reported in brackets (N = total number of the cells in all FOVs).

**Supplementary Figure 7**

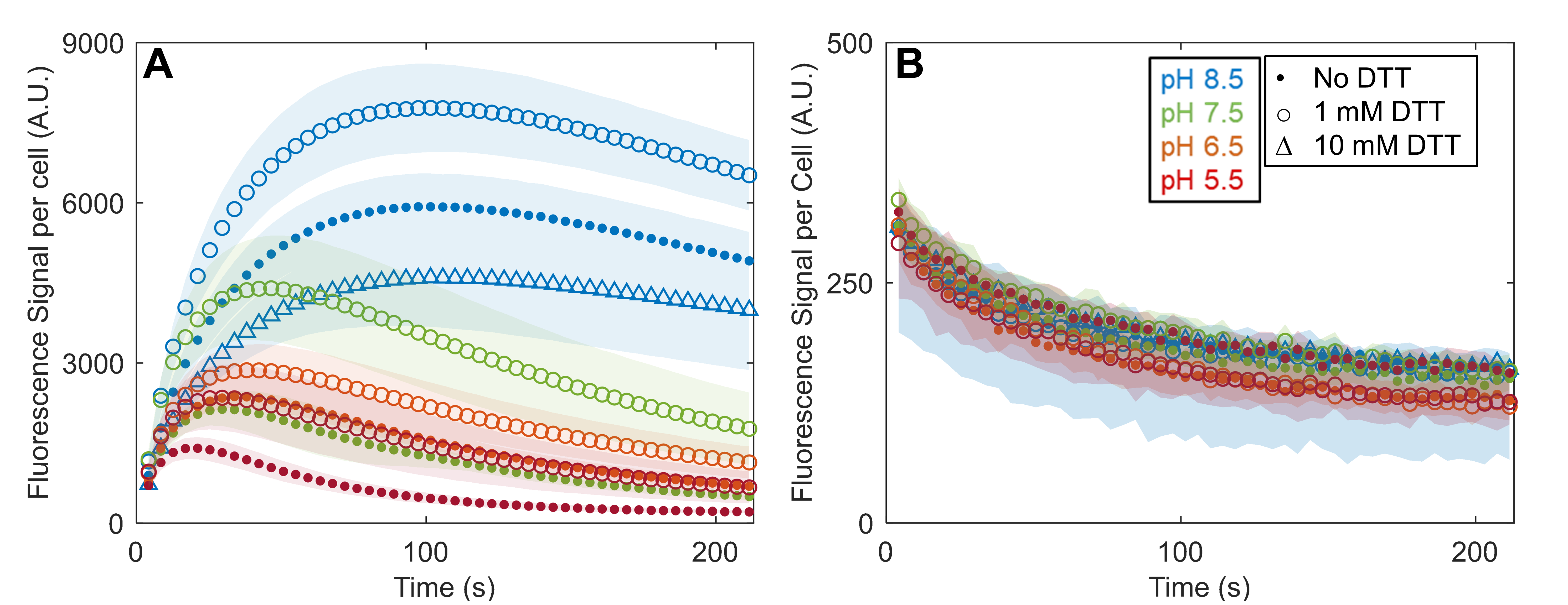

**Supplementary Figure 7: Effects of and DTT on mEos3.2 fluorescence signal in fixed *S. pombe* cells by point-scanning confocal microscopy.** Conditions: Time courses of fluorescence signal per *S. pombe* cell fixed with 2% formaldehyde in EMM5S medium for 30 min and imaged in various buffers under illumination at 405 nm (22 µW) and 561 nm (15 µW). Z-stacks with 19 slices from 8 FOVs of 85 µm x 85 µm were recorded at each time point. Plots are weighted mean (dots) and standard deviations (shaded area) of the fluorescence signal per cell. **(A)** *S. pombe* cells expressing cytoplasmic mEos3.2 under 9 different buffer conditions: (red dot) 50 mM MES (pH 5.5); (red circle) 50 mM MES (pH 5.5) with 1 mM DTT; (orange dot) 50 mM MES (pH 6.5); (orange circle) 50 mM MES (pH 6.5) with 1 mM DTT; (green dot) 50 mM Tris-HCl (pH 7.5); (green circle) 50 mM Tris-HCl (pH 7.5) with 1 mM DTT; (blue dot) 50 mM Tris-HCl (pH 8.5); (blue circle) 50 mM Tris-HCl (pH 8.5) with 1 mM DTT; (blue triangle) 50 mM Tris-HCl (pH 8.5) with 10 mM DTT. **(B)** Wild type *S. pombe* cells under the same 9 buffer conditions as panel A. Z-stacks with 19 slices from 4 FOVs of 85 µm x 85 µm were recorded at each time point and used for autofluorescence background subtraction.

**Supplementary Figure 8**

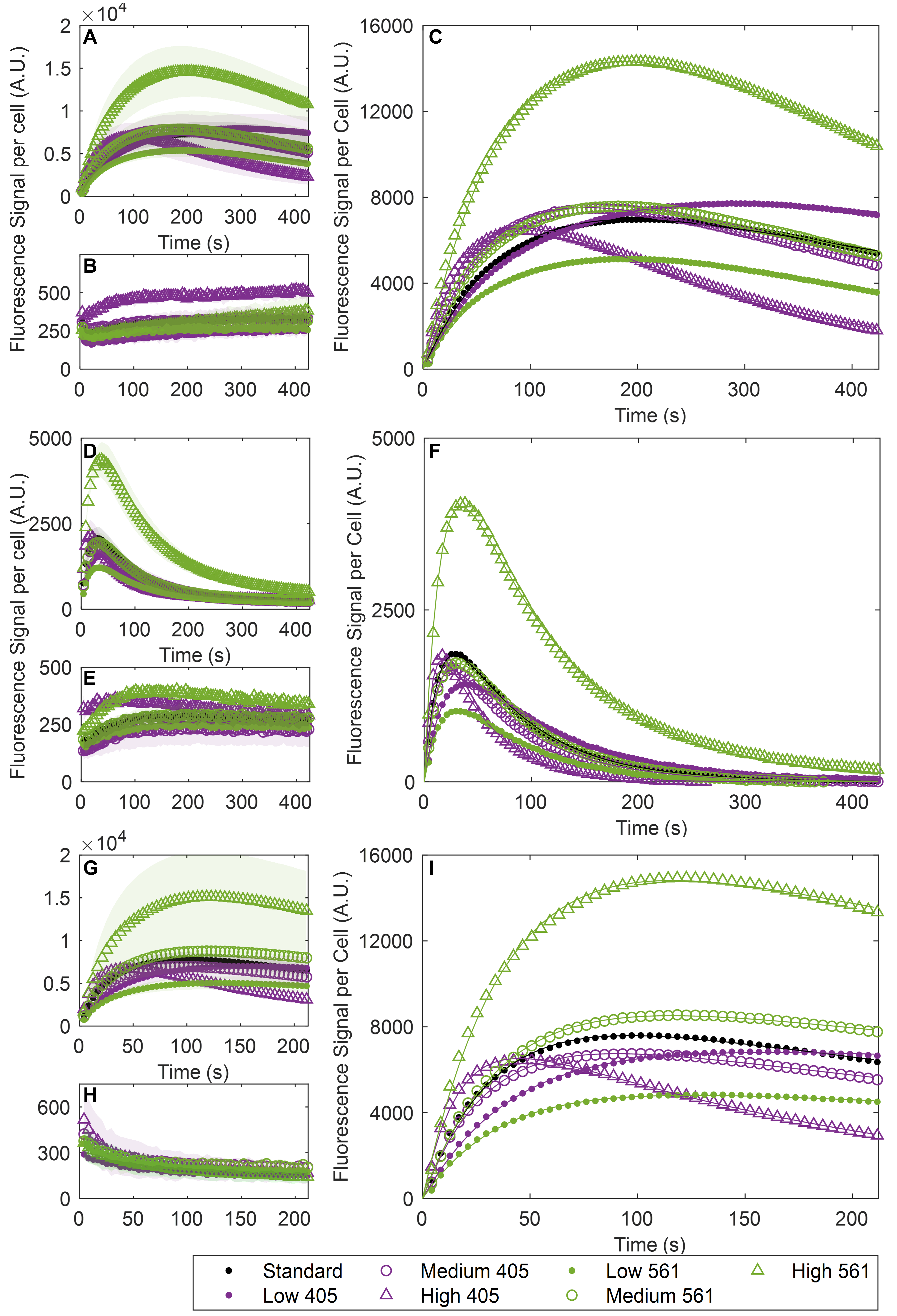

**Supplementary Figure 8: Effects of 405-nm and 561-nm intensities on mEos3.2 fluorescence signal in live and fixed *S. pombe* cells by point-scanning confocal microscopy. (A)** Time courses of the fluorescence signal per live cell expressing cytoplasmic mEos3.2 under 7 different laser intensities. Conditions: (black dot) standard conditions: 405 nm laser power at 22 µW, and 561 laser power at 15 µW; (purple dot) low 405 nm laser power at 16 µW; (purple circle) medium 405 laser power at 28 µW; (purple triangle) high 405nm laser power at 56 µW; (green dot) low 561 nm laser power at 11 µW; (green circle) medium 561 nm laser power at 19 µW; (green triangle) high 561 nm laser power at 37 µW. Z-stacks with 19 slices from 4 FOVs of 85 µm x 85 µm were recorded at each time point. Plots are weighted mean (dots) and standard deviation (shaded area) of the fluorescence signal per cell. **(B)** Time course of the autofluorescence signal per live wild type *S. pombe* cells under 7 different laser intensities. Z-stacks from two FOVs of 85 µm x 85 µm were recorded at each time point and used for autofluorescence background subtraction. Plots are weighted mean (dots) and standard deviation (shaded area) of the fluorescence signal per cell. **(C)** Eq. 7 of the 3-state model was fit to the time courses of mEos3.2 fluorescence signal per cell after autofluorescence background subtraction (dot or circle or triangle). The lines are theoretical curves with fitted parameters giving the best fit to the data. **(D-F)** *S. pombe* cells fixed with 2% formaldehyde in EMM5S medium for 30 min and imaged in EMM5S medium under the same 7 laser intensities as in Panel A. **(G-I)** *S. pombe* cells fixed with 2% formaldehyde for 30 min in EMM5S and imaged in 50 mM Tris-HCl buffer at pH 8.5 with 1 mM DTT under the same 7 laser intensities as in Panel A.

**Supplementary Figure 9**

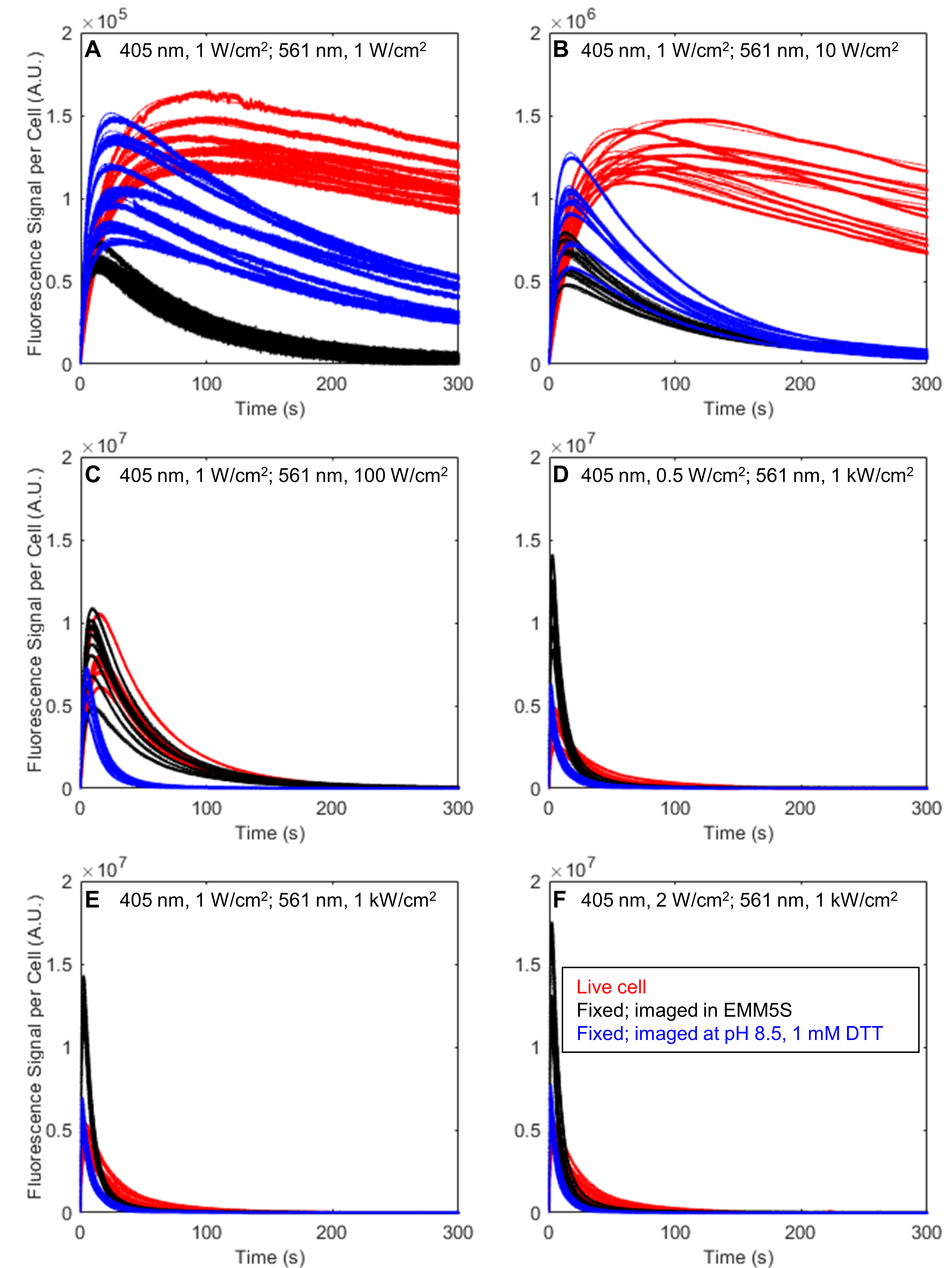

**Supplementary Figure 9: The effects of the 405-nm and 561-nm laser intensities on mEos3.2 fluorescence signal from live and fixed *S. pombe* cells by epi-fluorescence microscopy.** Time courses of the fluorescence signal per cell expressing cytoplasmic mEos3.2 in each FOV (after autofluorescence background subtraction): live cells (red dots) and cells fixed with 2% formaldehyde in EMM5S medium for 30 min and imaged in EMM5S (black dots) or imaged in 50 mM Tris-HCl buffer at pH 8.5 with 1 mM DTT (blue dots). Cells were illuminated continuously at 405 nm and 561 nm and imaged at 50 fps with 20-ms exposure time. The continuous lines are best fits of Eq. 7 of the 3-state model to the data (dots). Conditions: **(A)** 405 nm laser intensity at 1 W/cm^2^, 561 nm laser intensity at 1 W/cm^2^; **(B)** 405 nm laser intensity at 1 W/cm^2^, 561 nm laser intensity at 10 W/cm^2^; **(C)** 405 nm laser intensity at 1 W/cm^2^, 561 nm laser intensity at 100 W/cm^2^; **(D)** 405 nm laser intensity at 0.5 W/cm^2^, 561 nm laser intensity at 1 kW/cm^2^; **(E)** 405 nm laser intensity at 1 W/cm^2^, 561 nm laser intensity at 1 kW/cm^2^; **(F)** 405 nm laser intensity at 2 W/cm^2^, 561 nm laser intensity at 1 kW/cm^2^.

**Supplementary Table 1**

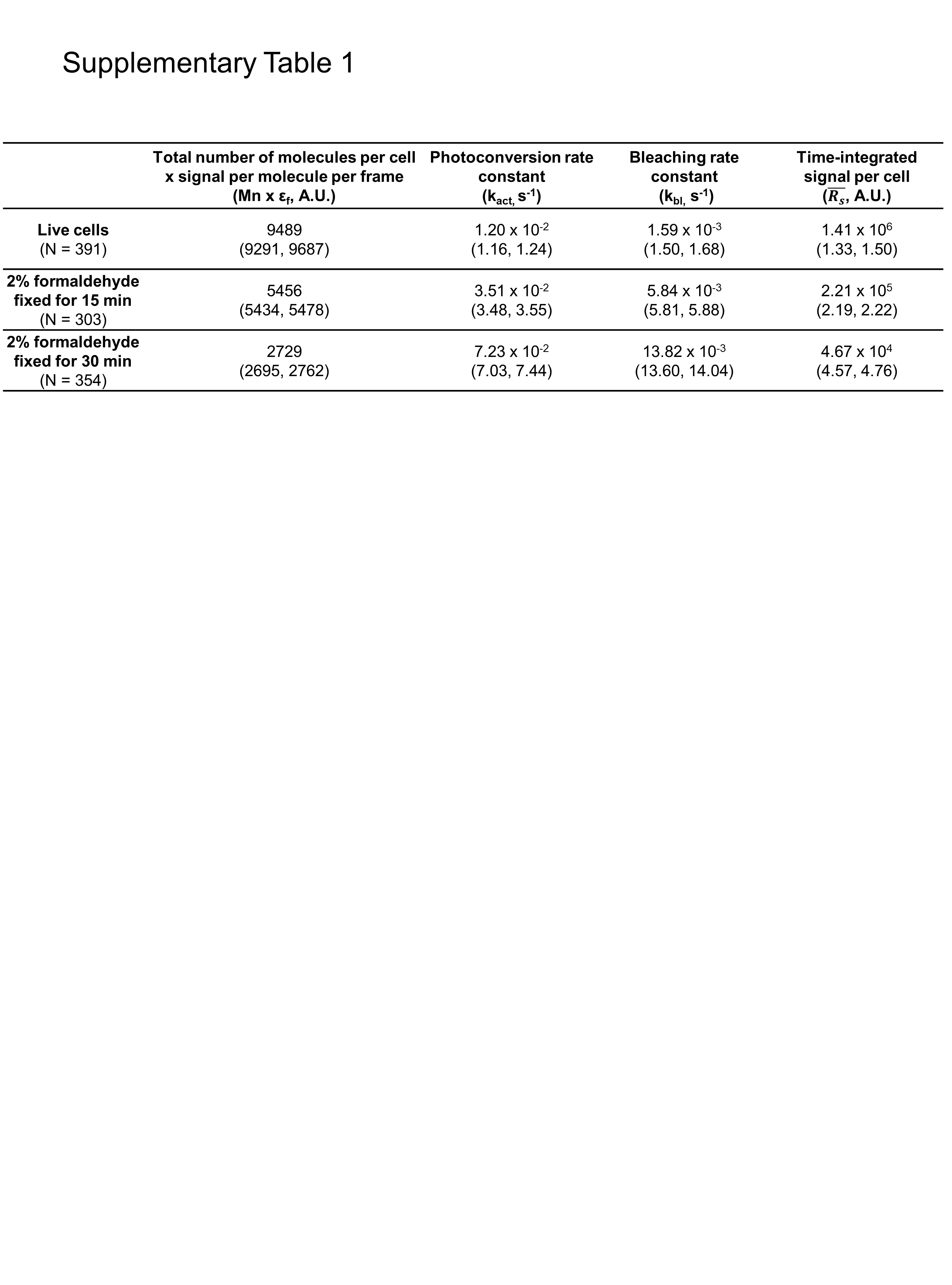

**Supplementary Table 1: Effects of fixation on the fluorescence signal and photoconversion and photobleaching rate constants of mEos3.2 in *S. pombe* cells measured by point-scanning confocal microscopy.** The table lists the product of the total number of molecules per cell and the signal of the R-state mEos3.2 molecule per frame (M_n_ x ε_f_), photoconversion rate constant (k_act_), and photobleaching rate constant (k_bl_) from fitting Eq. 7 of the 3-state model to the data, and the time-integrated signal per cell ($\bar{R_{s}}$) calculated from Eq. 8 (Fig. 2). The 95% confidence intervals are reported in the brackets. (N = total number of the cells in the FOVs)

**Supplementary Table 2**

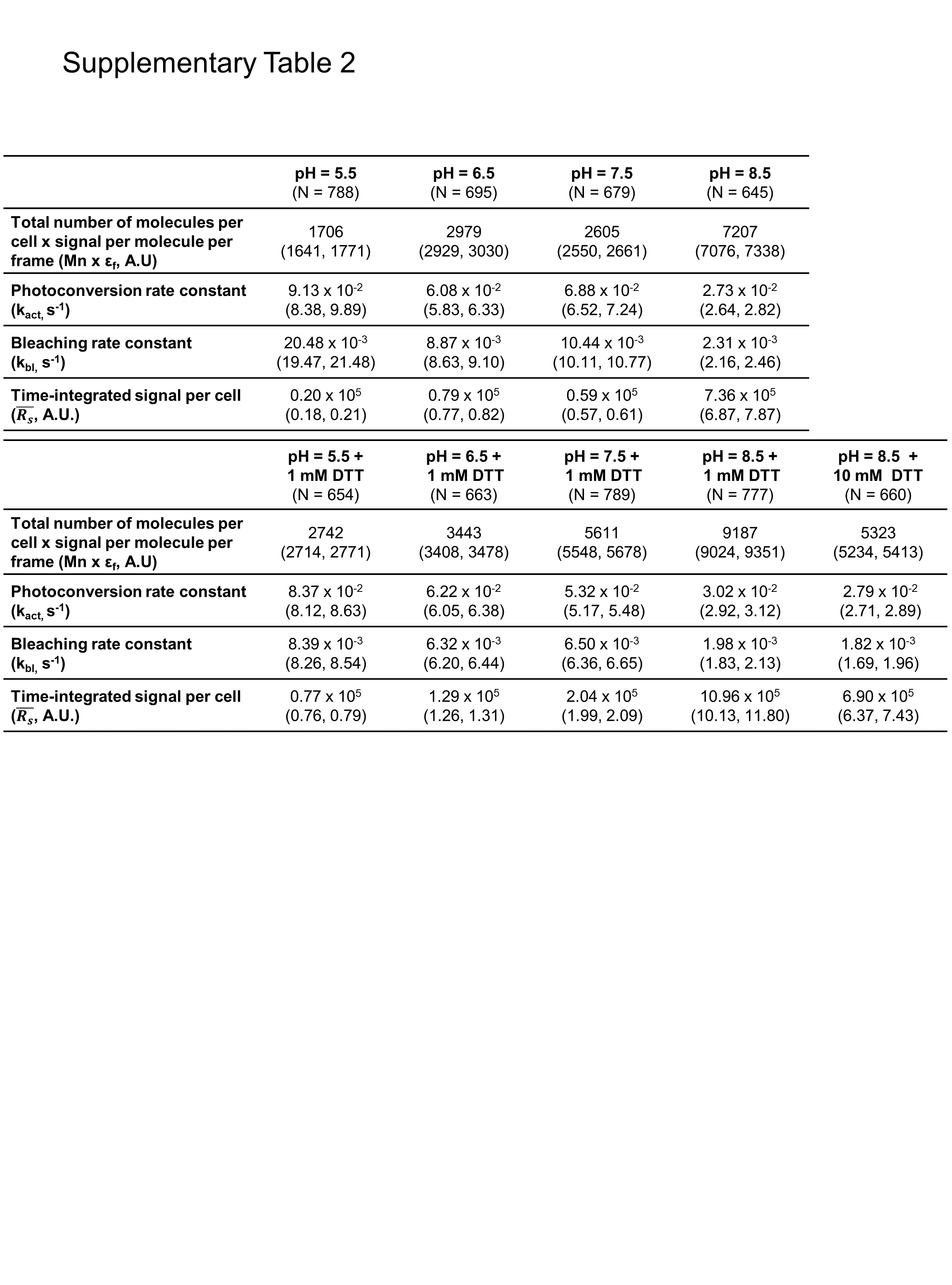

**Supplementary Table 2: Effects of pH and reducing agent on the fluorescence signal and photoconversion and photobleaching rate constants of mEos3.2 in fixed *S. pombe* cells measured by point-scanning confocal microscopy.** The table lists the product of the total number of molecules per cell and the signal the a R-state mEos3.2 molecule per frame (M_n_ x ε_f_), photoconversion rate constant (k_act_), and photobleaching rate constant (k_bl_) from fitting Eq. 7 of the 3-state model to the data, and the time-integrated signal per cell ($\bar{R_{s}}$) calculated from Eq. 8 (Fig. 3, S4). The 95% confidence intervals are reported in the brackets. (N = total number of the cells in the FOVs)

**Supplementary Table 3**

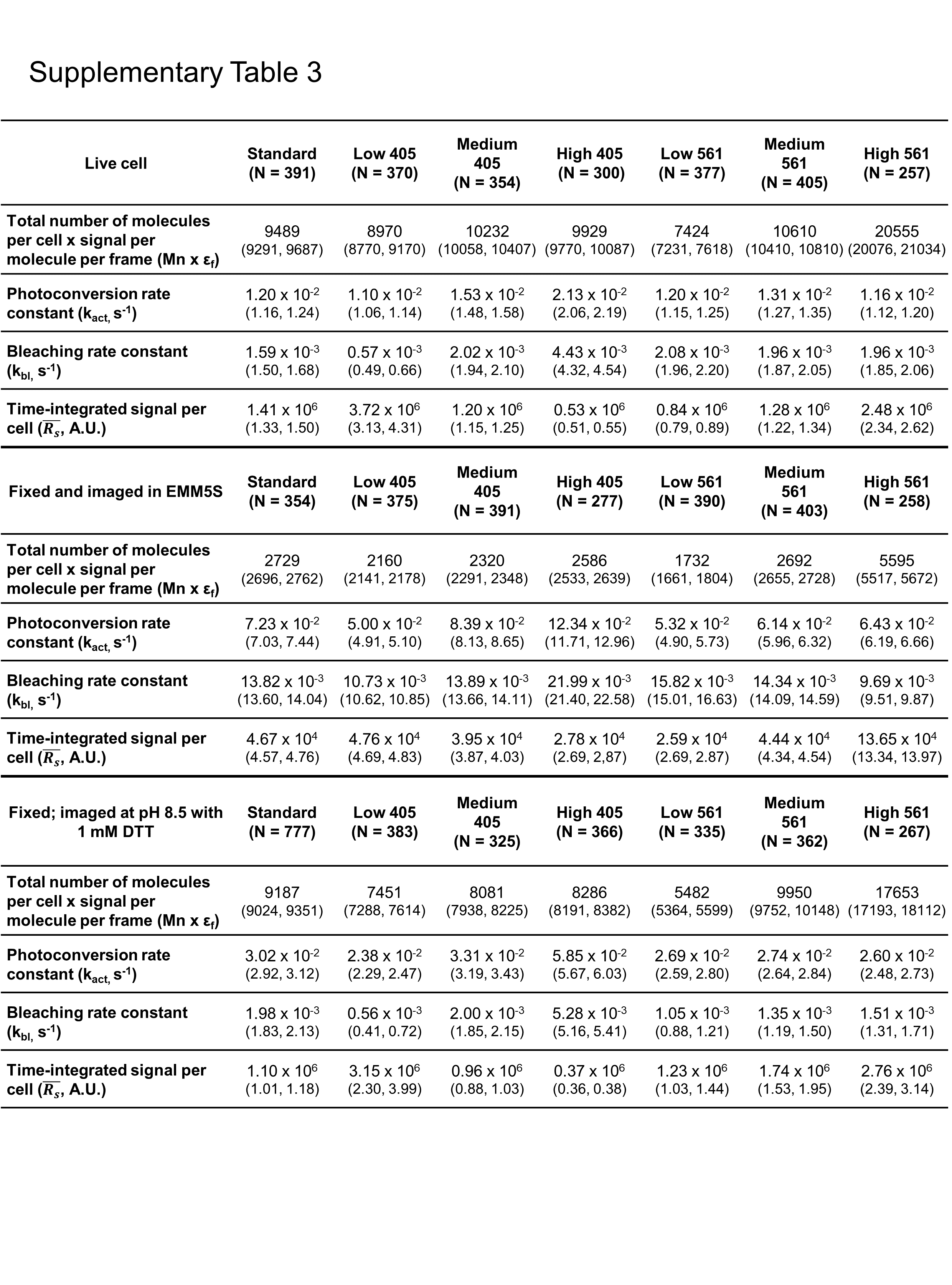

**Supplementary Table 3: Effects of laser intensities on the fluorescence signal and photoconversion and photobleaching rate constants of mEos3.2 in live and fixed *S. pombe* cells measured by point-scanning confocal microscopy.** The table lists the product of the total number of molecules per cell and the signal of the R-state mEos3.2 molecule per frame (M_n_ x ε_f_), photoconversion rate constant (k_act_), and photobleaching rate constant (k_bl_) from fitting Eq. 7 of the 3-state model to the data, and the time-integrated signal per cell ($\bar{R_{s}}$) calculated from Eq. 8 (Fig. 4, S5). The 95% confidence intervals are reported in the brackets. (N = total number of the cells in the FOVs)

**Supplementary Table 4**

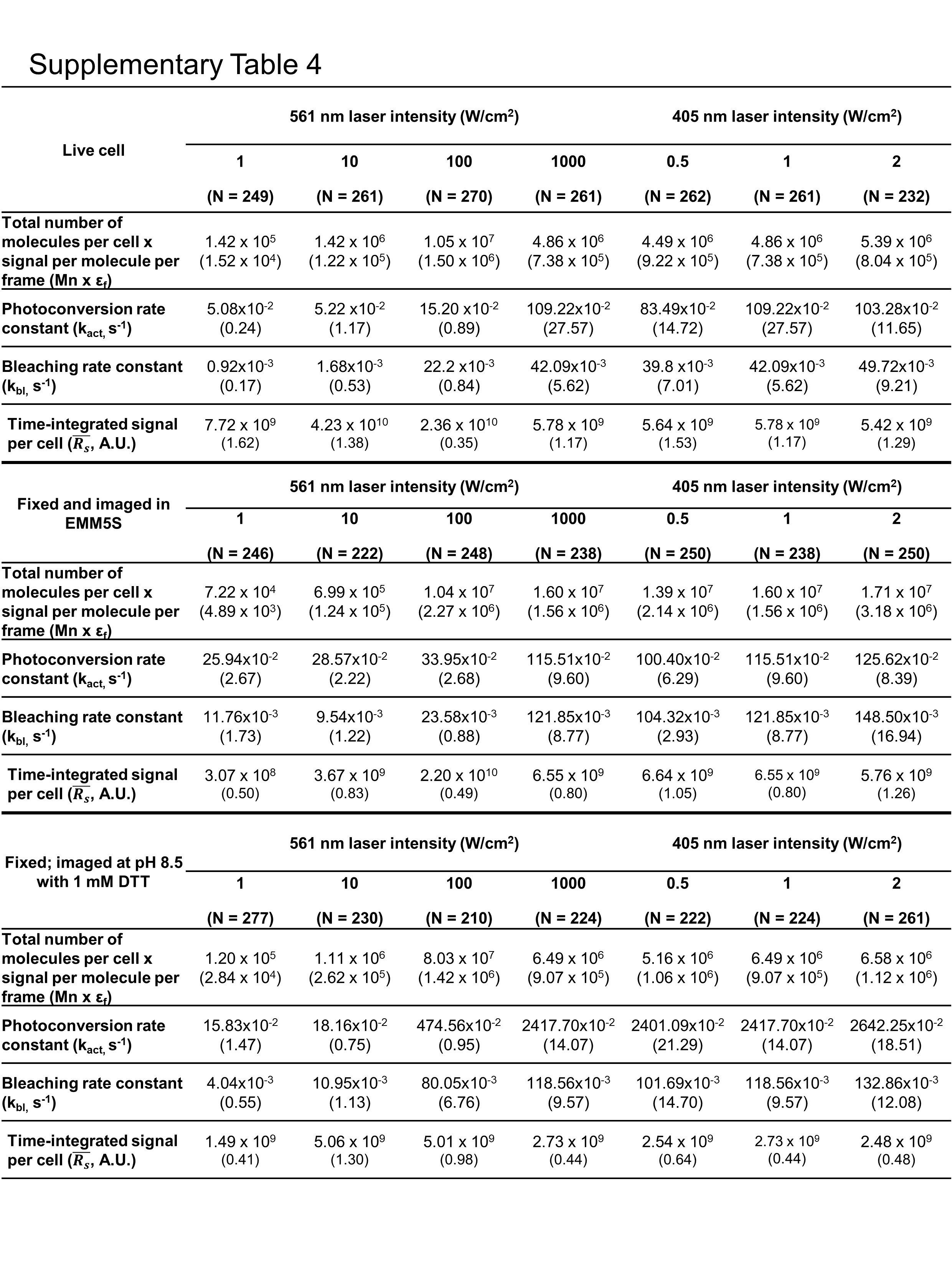

**Supplementary Table 4: Effects of laser intensities on the fluorescence signal and photoconversion and photobleaching rate constants of mEos3.2 in live and fixed *S. pombe* cells measured by widefield fluorescence microscopy.** The table lists the product of the total number of molecules per cell and the signal of the R-state mEos3.2 molecule per frame (M_n_ x ε_f_), photoconversion rate constant (k_act_), and photobleaching rate constant (k_bl_) from fitting Eq. 7 of the 3-state model to the data, and the time-integrated signal per cell ($\bar{R_{s}}$) calculated from Eq. 8 (Fig. 4, S6). The 95% confidence intervals are reported in the brackets. (N = total number of the cells in the FOVs)
